# Traumatic Brain Injury Associated with Altered Corpus Callosum Microstructure in Females: Exploring the Roles of Menopause Timing and Hormone Therapy in UK Biobank

**DOI:** 10.64898/2026.01.26.701743

**Authors:** Goretti España-Irla, Emma M. Tinney, Madeleine Perko, Mark Nwakamma, Stephanie Noble, Timothy P. Morris

## Abstract

**Background:** Traumatic brain injury (TBI) has lasting effects on white matter, yet sex-specific factors such as menopause timing and hormone replacement therapy (HRT) may modulate these outcomes in females.

**Objectives:** To investigate how TBI, menopause timing, HRT use, and reproductive history relate to corpus callosum white matter microstructure in female UK Biobank participants.

**Design:** Cross-sectional analysis of UK Biobank diffusion MRI data using propensity score matching to compare females with TBI to controls.

**Methods:** We analyzed diffusion MRI data from females with and without TBI. Corpus callosum fractional anisotropy (FA), mean diffusivity (MD), and isotropic volume fraction (ISOVF) were assessed. TBI effects were examined across pre- and post-menopausal groups, accounting for HRT use, duration, and reproductive factors.

**Results:** Females with TBI (n=363) exhibited widespread corpus callosum alterations compared to propensity-matched controls (n=10,128), with reduced FA across all regions (genu: β=-0.006, FDR p=0.027; body: β=-0.006, FDR p=0.002; splenium: β=-0.004, FDR p=0.009) and elevated MD in anterior regions (genu: FDR p=0.001; body: FDR p=0.002). TBI sustained before menopause was associated with significantly lower splenium FA (β=-0.010, p=0.031) and higher body MD (β=0.000019, p=0.021) compared with TBI sustained after menopause and controls. HRT use did not modify TBI-related alterations in primary analyses. However, among HRT users (n=3,108), a significant TBI×duration interaction emerged for genu MD (β=2.00×10^−6^, p=0.0295), indicating that the effect of HRT duration on white matter microstructure differed between TBI cases and healthy females. Reproductive factors (parity, reproductive lifespan) independently predicted some white matter measures but did not confound TBI, menopause timing, or HRT associations.

**Conclusions:** TBI-related white matter changes in females are influenced by menopause timing and hormonal exposure, with HRT effects dependent on duration and injury context. These findings highlight the importance of sex- and hormone-specific approaches in TBI research and the need for longitudinal studies to clarify mechanisms and potential interventions.

## Introduction

Women’s brain health remains an underexplored area of neuroscience and public health. Emerging evidence suggests that sex-specific biological processes, particularly reproductive transitions, influence brain structure and function with implications for cognitive aging and neurological disease (1). Menopause, defined as the permanent cessation of menstruation due to ovarian follicular depletion, typically occurs between ages 45 and 55 and is accompanied by marked declines in estrogen and progesterone (3). These hormonal changes have broad effects on the brain, as estrogen receptors are abundant in regions supporting cognition and emotion regulation (4–6). Neuroimaging studies demonstrate reductions in gray matter volume and disruptions in white matter microstructure during the menopausal transition (7). Diffusion tensor imaging (DTI) findings of decreased fractional anisotropy (FA) and increased mean diffusivity (MD) suggest compromised white matter integrity during this period of hormonal fluctuation (8– 12). Hormone replacement therapy (HRT) has been proposed as a means to counteract menopause-related neural changes. The “critical window hypothesis” posits that HRT may be neuroprotective when initiated near menopause onset, with some studies showing improved cognition and white matter integrity (13). However, results remain inconsistent, likely due to variation in formulation, dosage, duration, and individual factors (14–18).

Traumatic brain injury (TBI) represents another major threat to brain health, with growing recognition of sex differences in recovery outcomes (19). Some studies indicate that women experience longer recovery and more persistent symptoms after TBI compared to men, potentially due to interactions between injury mechanisms and hormonal status (19–25). The corpus callosum, the largest white matter tract, plays a key role in interhemispheric communication and is particularly vulnerable to diffuse axonal injury. DTI studies consistently show reduced FA and increased MD in this structure following TBI, correlating with cognitive deficits and functional impairment (26–28).

Although hormonal transitions and TBI independently alter white matter integrity, their interaction remains poorly understood. TBI disrupts axons and myelin, and long-term recovery involves remyelination processes that are sensitive to the hormonal milieu (29). Estrogen has been reported to support myelin repair and limit secondary injury (30–35), suggesting that menopause-related hormonal decline may reduce reparative capacity and increase vulnerability to chronic white matter degeneration. Consequently, the timing of TBI relative to menopause may influence both acute injury responses and long-term recovery trajectories. Injury sustained prior to menopause may occur in a hormonally supportive environment that facilitates repair, whereas injury sustained after menopause may unfold under diminished neuroprotection, or alternatively, pre-menopausal injury may interact with subsequent estrogen withdrawal to accelerate later-life white matter decline.

Together, these mechanisms indicate that menopausal status at the time of injury is a potential biologically meaningful modifier of long-term white matter integrity. Specifically, differences in hormonal environment at injury and during recovery may shape persistent alterations in diffusion-based measures of corpus callosum microstructure. The present study investigated corpus callosum microstructure in females who experienced TBI within ten years before neuroimaging, compared to matched controls. We aimed to: (1) assess TBI-related alterations in diffusion metrics (FA, MD, and Isotropic Volume Fraction (ISOVF)) in females; (2) determine whether the timing of TBI relative to menopause (i.e., TBI sustained before vs. after menopause) influences white matter integrity; and (3) evaluate whether HRT use moderates these associations.

## Methods

### Study Population and Design

This cross-sectional study was conducted and reported in accordance with the Strengthening the Reporting of Observational Studies in Epidemiology (STROBE) guidelines for cross-sectional studies. This study utilized data from the UK Biobank (UKB), a large prospective cohort of over 500,000 generally healthy adults aged 40-70 years at recruitment (2006-2010) and assessed at one of 22 centers across the United Kingdom. The first MRI scan commenced in 2014 and is planned to continue until approximately 100,000 participants are scanned at one of four assessment centers (Cheadle, Newcastle, Reading, and Bristol). Details on these assessments are available in open-access protocols. For the current analysis, we restricted our sample to female participants with available neuroimaging data from the imaging visit (Instance 2) and complete information on reproductive health variables including age at natural menopause. We excluded participants who had undergone hysterectomy or bilateral oophorectomy, as these surgical interventions would confound analyses of menopause timing and hormonal status. We further excluded participants with missing data on key variables including age at recruitment, socioeconomic status (Townsend deprivation index), educational attainment, cardiometabolic medication use (cholesterol-lowering medications, blood pressure medications, and insulin), HRT use, and age at the imaging visit.

### Ethical Approval

This secondary data analysis study was conducted under generic approval from the NHS National Research Ethics Service (approval letter dated 17th June 2011, ref. 11/NW/0382). Written informed consent was obtained from all participants in the study (consent for research, by UKB). Analyses were completed using UKB project-GxVJqV0JfzGZQKFgg95b97ZP.

### Traumatic Brain Injury Classification

History of TBI was ascertained using hospital episode statistics (HES) data with ICD-10 diagnostic codes. We employed both a narrow-band definition using 51 ICD-10 codes specific to TBI and associated terms, and a broad-band definition using a comprehensive list of 1,798 relevant ICD-10 codes capturing data on head injury (Supplementary Table 1). Primary and secondary ICD-10 diagnoses were based on HES admission NHS data, as detailed by the open-access protocol (https://biobank.ctsu.ox.ac.uk/crystal/crystal/docs/HospitalEpisodeStatistics.pdf), from May 1995 to October 2021 and as applied in previous published studies (27). Participants who had a relevant narrow-band or broad-band ICD code at any date prior to the imaging visit were included as TBI cases; a narrow-band case was necessarily also a broad-band case. For the primary analyses, we combined narrow-band and broad-band TBI cases into a single TBI exposure group. To examine the relatively recent effects of brain injury while maintaining adequate sample size, we restricted TBI cases to those with injury occurring 0-10 years prior to the imaging visit (Instance 2). Time since injury was calculated as the difference between the date of the first TBI-related hospital admission and the date of the imaging assessment. Among TBI cases, we further classified participants according to menopause timing relative to injury: pre-menopause TBI (TBI sustained before natural menopause) versus post-menopause TBI (TBI sustained after natural menopause).

### Neuroimaging Outcomes

MR 3T image sequence, equipment, acquisition and processing details are available open-access on the UKB website in the form of protocol (http://biobank.ctsu.ox.ac.uk/crystal/refer.cgi?id=2367) and imaging documentation (http://biobank.ctsu.ox.ac.uk/crystal/refer.cgi?id=1977). Detailed methodologies are described in previous open-access reports. For imaging data, we used imaging-derived phenotypes processed (including quality control) by UKB. The selection of brain regions and DTI metrics analyzed was guided by prior literature (27), ensuring focus on measures relevant to TBI-related microstructural changes. Specifically, we accessed diffusion tensor imaging (DTI) metrics including fractional anisotropy (FA), mean diffusivity (MD), and isotropic volume fraction (ISOVF) within the corpus callosum. Higher FA values reflect greater white matter integrity and more organized fiber tracts, while higher MD and ISOVF values indicate tissue pathology and reduced microstructural coherence. We analyzed three sub-regions of the corpus callosum (genu, body, and splenium) across these three DTI metrics, yielding a total of nine corpus callosum measures.

### Statistical Analysis

All analyses were performed using R version 4.4.0. Control participants were selected from female UKB participants with no history of TBI according to either narrow-band or broad-band definitions, meeting the same inclusion and exclusion criteria as TBI cases. To minimize confounding and ensure comparability between TBI and control groups, we employed full matching using the *MatchIt* package in R. Full matching creates matched sets where each TBI case is matched to one or more controls and each control may be matched to one or more TBI cases, thereby utilizing the entire eligible sample while optimizing covariate balance. Propensity scores were estimated using logistic regression with TBI status as the outcome and the following covariates as predictors: age at imaging visit (Instance 2), Townsend deprivation index at recruitment, highest educational qualification, and use of cardiometabolic medications. We used a caliper of 0.2 standard deviations of the propensity score logit to ensure adequate match quality. The matching approach targeted the average treatment effect on the treated (ATT), focusing inference on the TBI population. Full matching produces optimal weights for each participant that were subsequently incorporated into all outcome analyses. Covariate balance after matching was assessed using standardized mean differences, with values less than 0.10 considered indicative of adequate balance. Balance diagnostics were visualized using Love plots generated with the *cobalt* package (Supplementary Figure 1). The final matched sample comprised 363 TBI cases and 10,115 controls.

We utilized robust weighted linear regression models to estimate associations between TBI and corpus callosum microstructure. This analytical approach combines propensity score weighting with covariate adjustment to provide robust effect estimates even if either the propensity score model or outcome model is misspecified. We report standardized betas (β) on the per-standard deviation scale based on linear regressions for continuous outcomes, plus 95% confidence intervals. All models controlled for age at time of imaging, Townsend deprivation index, educational attainment, and cardiometabolic medication use. Among TBI cases only, we compared corpus callosum outcomes between pre-menopause and post-menopause TBI groups using weighted linear regression with adjustment for the aforementioned covariates. To examine whether HRT use modified the association between TBI and brain outcomes, we tested TBI×HRT interaction terms using analysis of variance (ANOVA) in models that included both main effects and their interaction, weighted by propensity scores. We corrected for multiple testing using false discovery rate (FDR) correction across all corpus callosum regions within each DTI metric (FA, MD, ISOVF). Statistical significance was set at FDR-corrected P < 0.05.

To assess the robustness of our findings, we conducted several sensitivity analyses. First, we examined whether reproductive factors (reproductive lifespan calculated as age at menopause minus age at menarche, and parity defined as number of live births) confounded the association between TBI and corpus callosum outcomes. We compared models with and without these reproductive factors to assess whether inclusion substantially altered TBI effect estimates (defined as >20% change in beta coefficients). Second, among females who reported ever using HRT, we examined whether HRT duration (calculated as age stopped minus age started for past users, or current age minus age started for current users) and age at HRT initiation were associated with corpus callosum outcomes. We tested whether these HRT characteristics predicted white matter microstructure independently and whether their effects differed by TBI status through interaction terms (TBI×duration and TBI×age initiated). Cases with implausible HRT duration data (negative values indicating stopped before started, or durations >50 years) were excluded from the analyses.

## Results

### Sample Characteristics

The final analytical sample comprised 10,491 female participants, including 363 females with a history of TBI occurring 0-10 years prior to neuroimaging and 10,128 matched controls without TBI history. Demographic and clinical characteristics are displayed in Table 1.

**Table 1.** Values are presented as mean (SD) for continuous variables and n (%) for categorical variables. Abbreviations: TBI: traumatic brain injury; SD: standard deviation; MRI: magnetic resonance imaging; BMI: body mass index; HRT: hormone replacement therapy.

| Variable | Overall<br>N = 10491 | Control<br>N = 10128 | TBI<br>N = 363 |
| --- | --- | --- | --- |
| Age at MRI scan, years | 65.51 (7.14) | 65.45 (7.11) | 67.18 (7.71) |
| Age at first TBI, years | 62.31 (8.22) | NA | 62.31 (8.22) |
| Age at recruitment, years | 54.24 (7.01) | 54.19 (6.99) | 55.47 (7.28) |
| Age at menopause, years | 50.93 (4.04) | 50.93 (4.02) | 50.84 (4.60) |
| Years of education | 13.39 (2.90) | 13.39 (2.90) | 13.41 (2.97) |
| Townsend deprivation index | -1.83 (2.74) | -1.83 (2.74) | -1.85 (2.77) |
| BMI, kg/m<sup>2</sup> | 25.99 (4.83) | 26.00 (4.83) | 25.83 (4.79) |
| Waist circumference, cm | 83.49 (12.00) | 83.47 (11.98) | 83.96 (12.34) |
| Cardiometabolic medications | 2,943 (28%) | 2,829 (28%) | 114 (31%) |
| Ever used HRT | 3,761 (36%) | 3,608 (36%) | 153 (42%) |
| Number of live births | 1.73 (1.17) | 1.73 (1.17) | 1.64 (1.20) |
| Reproductive span, years | 37.93 (4.34) | 37.93 (4.32) | 37.81 (4.80) |

TBI-Related Alterations in Corpus Callosum Microstructure Females with a history of TBI exhibited widespread alterations in corpus callosum white matter microstructure compared to matched controls (Table 2). Fractional anisotropy (FA), a marker of white matter integrity, was significantly reduced across all three corpus callosum sub-regions in TBI cases, with the largest reductions observed in the body and genu (both FDR p < 0.03). Mean diffusivity (MD), an indicator of tissue pathology, was significantly elevated in the genu and body of the corpus callosum among TBI cases compared to controls (FDR p = 0.001 and p = 0.002, respectively). Isotropic volume fraction (ISOVF), reflecting free water content and tissue disorganization, was significantly elevated in the genu (FDR p = 0.015) among TBI cases. However, ISOVF differences in the body and splenium did not reach statistical significance after FDR correction.

**Table 2.** Values represent standardized regression coefficients (β) from doubly robust weighted linear models adjusted for age at imaging, Townsend deprivation index, educational attainment, and cardiometabolic medication use. Abbreviations: TBI: traumatic brain injury; FA: fractional anisotropy; MD: mean diffusivity; ISOVF: isotropic volume fraction; FDR: false discovery rate; CI: confidence interval.

| Variable | Coefficient ($\beta$) | P-value | FDR P-value | Lower 95% CI | Upper 95% CI | R-squared | n (TBI) | n(Control) |
| --- | --- | --- | --- | --- | --- | --- | --- | --- |
| Genu FA | -0.006 | 0.001 | **0.002** | -0.010 | -0.002 | 0.214 | 284 | 8369 |
| Body FA | -0.006 | <0.001 | **0.002** | -0.009 | -0.002 | 0.163 | 284 | 8369 |
| Splenium FA | -0.004 | 0.004 | **0.009** | -0.006 | -0.001 | 0.047 | 284 | 8369 |
| Genu MD | 0.000 | <0.001 | **0.001** | 0.000 | 0.000 | 0.263 | 284 | 8369 |
| Body MD | 0.000 | <0.001 | **0.002** | 0.000 | 0.000 | 0.222 | 284 | 8369 |
| Splenium MD | 0.000 | 0.043 | 0.056 | 0.000 | 0.000 | 0.118 | 284 | 8369 |
| Genu ISOVF | 0.003 | 0.010 | **0.015** | 0.001 | 0.006 | 0.173 | 284 | 8369 |
| Body ISOVF | 0.002 | 0.167 | 0.188 | -0.001 | 0.004 | 0.224 | 284 | 8369 |
| Splenium ISOVF | 0.000 | 0.936 | 0.936 | -0.002 | 0.002 | 0.055 | 284 | 8369 |

### Menopause Timing and TBI-Related White Matter Alterations

Among the TBI cases, 30 females sustained their injury before menopause and 333 after menopause. Weighted ANOVA comparing healthy controls (n=10,128), pre-menopause TBI (n=30), and post-menopause TBI (n=333) revealed significant group differences across multiple corpus callosum measures (Supplementary Table 2, Part A). Significant group differences were observed for FA in all corpus callosum sub-regions: body (F=6.411, p=0.0016, FDR p=0.0019), genu (F=8.226, p<0.001, FDR p<0.001), and splenium (F=7.161, p<0.001, FDR p=0.0013). MD showed significant group differences in the body (F=9.9996, p<0.0001, FDR p<0.0001) and genu (F=8.956, p<0.0001, FDR p<0.0001). ISOVF in the genu also demonstrated significant group differences (F=4.031, p=0.0178, FDR p=0.0178).

Post-hoc pairwise comparisons (Supplementary Table 3, Part A and Figure 2) revealed that for body FA, both TBI groups showed reduced values compared to controls (pre-menopause: β=0.017, p=0.0034; post-menopause: β=0.004, p=0.0367). For body MD, all three groups differed significantly from each other (controls vs pre-menopause: β=-0.000024, p=0.0007; controls vs post-menopause: β=-0.0000057, p=0.0277; pre-menopause vs post-menopause: β=0.000019, p=0.0210). In the genu, both TBI groups showed reduced FA compared to controls (pre-menopause: β=0.015, p=0.0338; post-menopause: β=0.005, p=0.0254) and elevated MD (pre-menopause: β=-0.000023, p=0.0072; post-menopause: β=-0.0000079, p=0.0078). For splenium FA, pre-menopause TBI cases showed significantly lower values than both controls (β=0.013, p=0.0028) and post-menopause TBI cases (β=-0.010, p=0.0311), while post-menopause TBI did not differ significantly from controls (β=0.0025, p=0.1447) (Figure 2).

**Figure 1.**
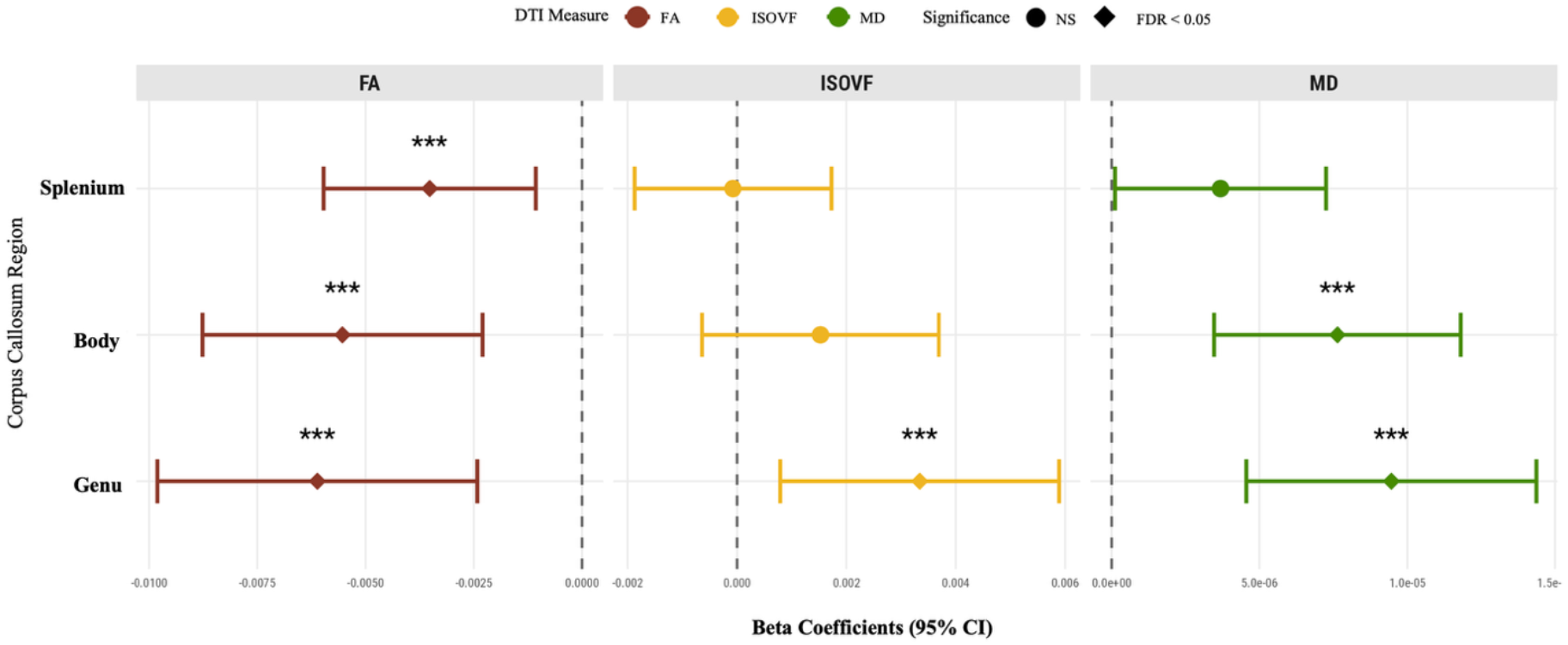
Corpus callosum microstructure by TBI status. Forest plot showing standardized regression coefficients (β) and 95% CIs for DTI metrics across corpus callosum sub-regions. Filled diamonds (***) indicate FDR p < 0.05. Dashed vertical lines represent no difference between groups; individuals with history of TBI and matched controls without history of TBI. Models adjusted for age at imaging, Townsend deprivation index, educational attainment, and cardiometabolic medication use. Abbreviations: TBI: traumatic brain injury; FA: fractional anisotropy; MD: mean diffusivity; ISOVF: isotropic volume fraction; FDR: false discovery rate; CI: confidence interval; NS: not significant.

**Figure 2.**
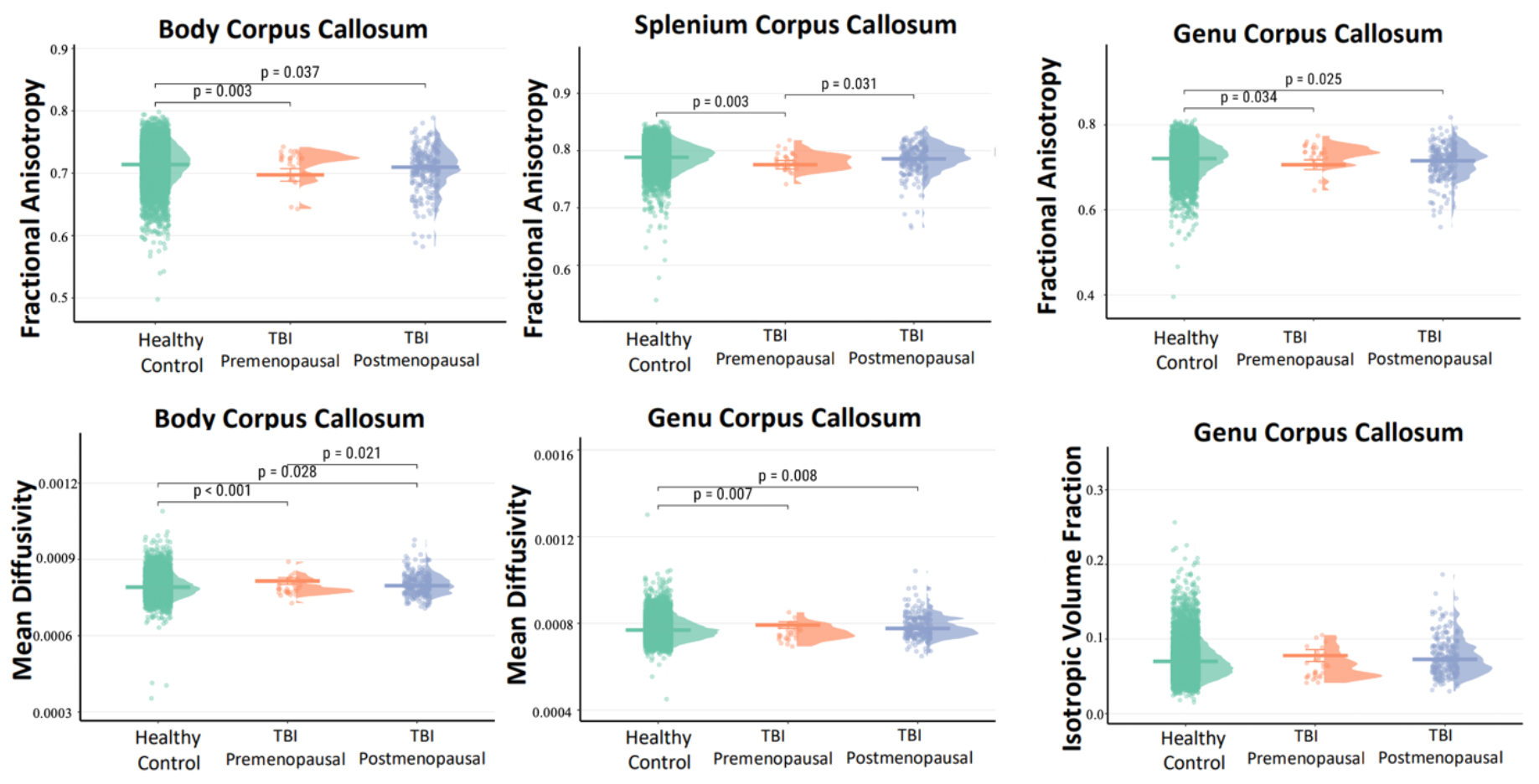
Corpus callosum white matter integrity by TBI status and menopause timing at injury. Raincloud plots showing the distribution of fractional anisotropy values across corpus callosum sub-regions (body, splenium, and genu) stratified by TBI status and menopause timing at injury. Horizontal lines represent median values. P-values indicate pairwise group comparisons. Models adjusted for age at imaging, Townsend deprivation index, educational attainment, and cardiometabolic medication use. Abbreviations: TBI: traumatic brain injury.

### Hormone Replacement Therapy and White Matter Microstructure

Among postmenopausal females, the sample comprised 6,436 healthy controls without HRT, 3,572 healthy controls with HRT, 187 TBI cases without HRT, and 146 TBI cases with HRT. Weighted ANOVA revealed significant group differences across multiple corpus callosum measures (Table S2, Part B). Significant group differences were observed for FA in the body (F=8.226, p<0.001, FDR p<0.001), genu (F=6.411, p=0.0003, FDR p<0.001), and splenium (F=7.161, p<0.001, FDR p<0.001). MD demonstrated significant group differences in the genu (F=28.041, p<0.0001, FDR p<0.0001) and body (F=14.436, p<0.0001, FDR p<0.0001). ISOVF in the genu showed significant group differences (F=28.041, p<0.0001, FDR p<0.0001).

Post-hoc pairwise comparisons (Supplementary Table 3, Part B, and Figure 3) revealed consistent differences between healthy females with and without HRT across all significant measures. Healthy HRT users showed reduced FA in the body (β=0.0033, p<0.0001) and genu (β=0.0041, p<0.0001), elevated MD in the body (β=-0.0000051, p<0.0001) and genu (β=-0.0000084, p<0.0001), and elevated ISOVF in the genu (β=-0.0044, p<0.0001) compared to healthy non-users. Comparisons between healthy controls without HRT and TBI groups showed that TBI without HRT differed significantly in genu FA (β=0.0072, p=0.0398) and genu MD (β=-0.000010, p=0.0228). TBI with HRT differed from healthy controls without HRT in body MD (β=-0.0000093, p=0.0230) and genu MD (β=-0.000012, p=0.0089). Direct comparisons between TBI groups (with versus without HRT) revealed no significant differences for any measure (all p>0.05).

**Figure 3.**
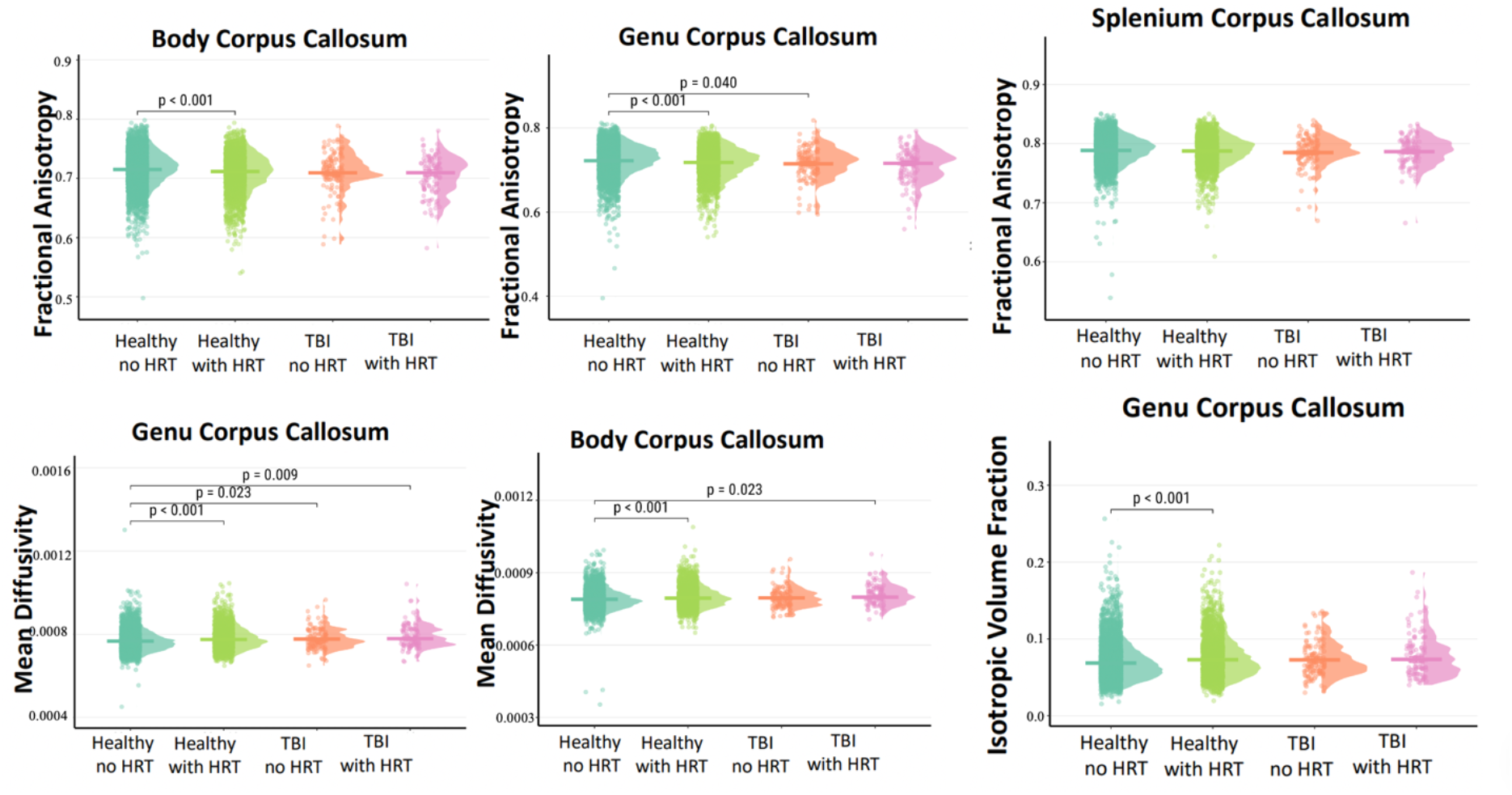
Corpus callosum white matter integrity by TBI status and HRT use in postmenopausal females. Raincloud plots showing the distribution of fractional anisotropy (top row), mean diffusivity (bottom left and center panels), and isotropic volume fraction (bottom right panel) across corpus callosum sub-regions (body, genu, and splenium) stratified by TBI status and hormone replacement therapy use. Horizontal lines represent median values. P-values indicate pairwise group comparisons. Models adjusted for age at imaging, Townsend deprivation index, educational attainment, and cardiometabolic medication use. Abbreviations: TBI: traumatic brain injury; HRT: hormone replacement therapy.

### Sensitivity Analyses

#### Propensity Score Matching Quality

Assessment of propensity score overlap revealed good common support between TBI cases and controls, with a common support region of [0.0189, 0.0604]. Only 17 control participants (0.2%) fell outside this region, with all TBI cases within the common support region. Propensity score distributions showed substantial overlap between groups, and matching weights were well-distributed, with 141 controls (1.4%) receiving extreme weights (>10). These diagnostics confirm adequate covariate balance and overlap, supporting the validity of the propensity score weighting approach.

#### Reproductive Factors Analysis

To assess whether reproductive factors confounded the observed TBI effects, we reanalyzed all significant corpus callosum outcomes including reproductive lifespan and parity as additional covariates. The inclusion of these reproductive factors produced minimal changes in TBI effect estimates, with percent changes in beta coefficients ranging from -0.4% to -8.7%. All TBI associations remained statistically significant after adjustment, with effect sizes and confidence intervals nearly identical to the primary analysis. Both reproductive factors independently predicted several corpus callosum measures: parity was associated with genu FA (β=0.00087, p=0.002), genu MD (β=-0.00000, p<0.001), body MD (β=-0.00000, p<0.001), genu ISOVF (β=-0.00111, p<0.001), and splenium FA (β=-0.00121, p<0.001). Reproductive lifespan was associated with genu FA (β=0.00017, p=0.037). However, despite these independent associations with white matter microstructure, inclusion of reproductive factors did not substantially alter TBI effect estimates, indicating that reproductive history does not confound the relationship between TBI and corpus callosum integrity in this sample (Supplementary Table 4).

#### HRT Duration and Timing Analysis

Among HRT users with valid duration data (n=3,108), we examined whether HRT duration or age at initiation influenced corpus callosum outcomes. HRT duration showed a significant association with genu MD (β=-0.000000, p=0.0145) and splenium FA (β=0.000530, p<0.0001), with longer duration associated with higher MD and higher FA respectively. A significant TBI×duration interaction was observed for genu MD (β=2.00×10^−6^, p=0.0295) (Figure 4), suggesting that the effect of HRT duration on this measure differed between TBI cases and controls. Age at HRT initiation was significantly associated with genu ISOVF (β=-0.000270, p=0.0049), with earlier initiation associated with higher ISOVF values. However, the majority of corpus callosum measures showed no significant associations with either HRT duration or age at initiation (all other p>0.05). These findings suggest limited and region-specific effects of HRT characteristics on white matter microstructure within the range observed in this sample (Supplementary Table 5).

**Figure 4.**
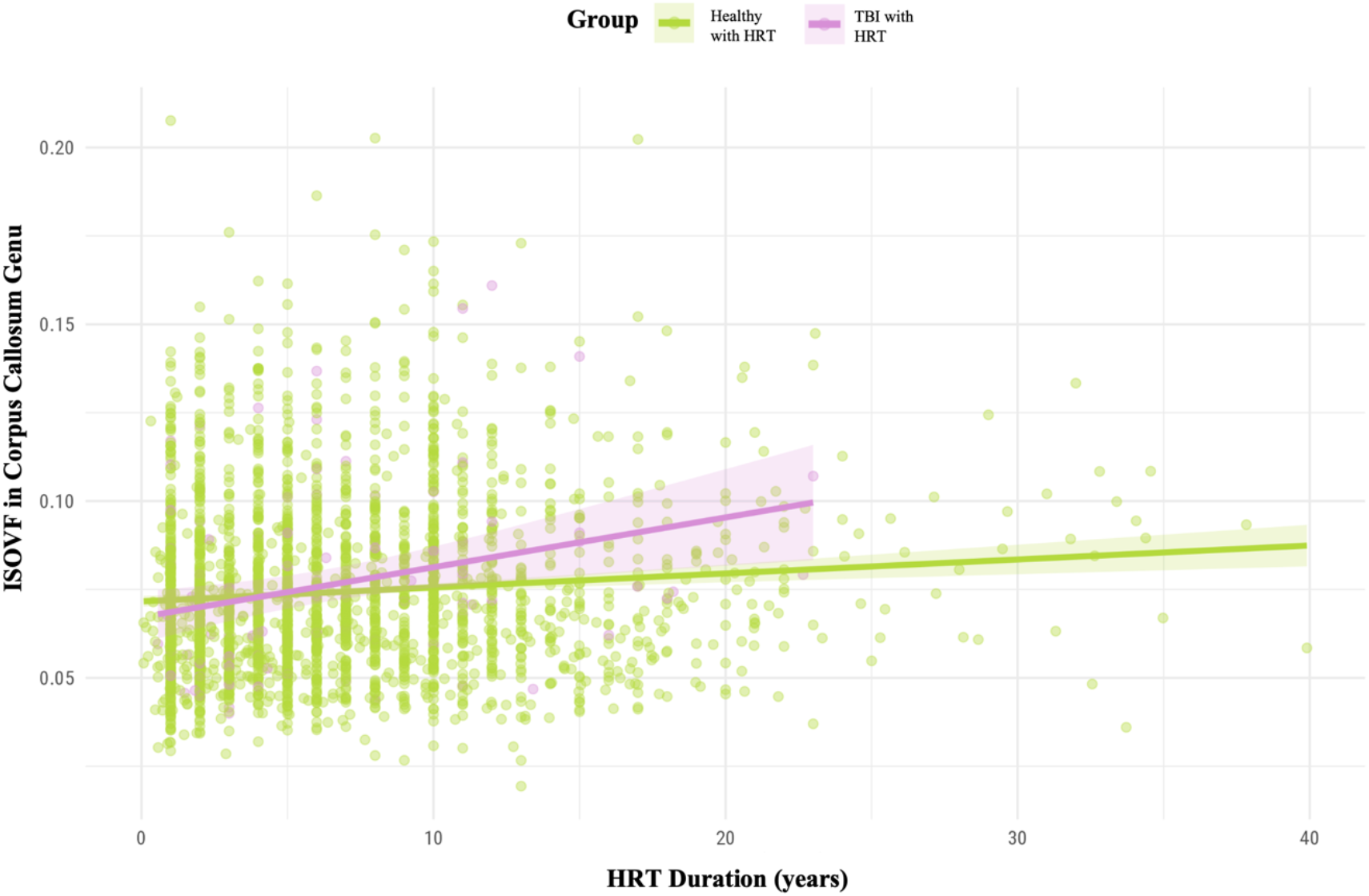
Interaction between hormone replacement therapy duration and TBI status on genu mean diffusivity. Scatter plot showing the relationship between HRT duration (years) and mean diffusivity in the genu corpus callosum for postmenopausal females with and without TBI history. Individual data points are shown for healthy females with HRT (green) and TBI females with HRT (purple), with fitted regression lines for each group. Models adjusted for age at imaging, Townsend deprivation index, educational attainment, and cardiometabolic medication use. Abbreviations: HRT: hormone replacement therapy; TBI: traumatic brain injury; MD: mean diffusivity.

## Discussion

This study examined the relationship between TBI, menopause timing, and HRT use on corpus callosum white matter microstructure in female UK Biobank participants. Building on prior research demonstrating these alterations across both sexes (27), females with a TBI history in the last 10 years exhibited widespread corpus callosum alterations compared to matched controls, with reduced FA and elevated MD and ISOVF across multiple sub-regions. Effects were most pronounced in the anterior corpus callosum, particularly the genu, aligning with literature showing greater age- and injury-related vulnerability in anterior corpus callosum subregions (36,37).

Our analysis of menopause timing revealed that pre-menopausal TBI cases showed lower splenial FA and higher MD in the corpus callosum body compared with both controls and post-menopausal TBI cases, whereas post-menopausal TBI cases did not differ from controls in splenial FA. The mean age at first TBI in our pre-menopausal group was 49.03 years, indicating that most of these cases were potentially perimenopausal at the time of injury. This aligns with prior literature suggesting a window of heightened vulnerability in the years preceding menopause, when hormonal fluctuations and increased inflammatory activity may amplify brain susceptibility and thereby exacerbate TBI-related microstructural alterations (38,39). Reproductive history, including parity and reproductive lifespan, was independently associated with white matter microstructure but did not substantially alter TBI effect estimates, indicating that lifetime reproductive factors do not explain the observed differences between pre- and post-menopausal TBI groups. Together, these findings suggest that menopause timing itself, or the concurrent hormonal environment, may modulate vulnerability to TBI.

Investigation of HRT revealed that, among healthy postmenopausal females, users exhibited lower FA and higher MD compared with non-users, consistent with recent evidence that HRT effects on brain structure are context-dependent and not uniformly neuroprotective (15– 18,40–42), challenging earlier assumptions about universal neuroprotection (43–47). Importantly, HRT did not significantly modify TBI-related white-matter differences, both TBI groups displayed similar patterns independent of HRT status. This suggests that, in this observational cohort, HRT was not associated with measurable improvement in existing structural changes, though its influence cannot be interpreted as definitively beneficial or detrimental. However, effects could depend on under-characterized factors such as formulation, dosage, timing, duration, or the interval between injury and treatment. Additionally, emerging evidence highlights the importance of route of administration, specific estrogen and progestin composition, and genetic factors (13– 15,48–50), suggesting that future work should account for these variables when assessing potential HRT–TBI interactions.

Since our primary analyses relied on a binary indicator of HRT use, we examined whether more nuanced characteristics, specifically timing of initiation and duration, might influence these patterns, consistent with recent research showing that these specifications can meaningfully affect brain aging trajectories in females (14). Among our healthy female group, white matter integrity remained largely stable across varying HRT durations, indicating that extended hormone use alone may not substantially alter corpus callosum structure. In contrast, postmenopausal females who sustained a TBI and used HRT showed increasing genu MD with longer HRT duration, suggesting that prolonged hormone exposure may interact with postmenopausal brain vulnerability to influence white matter outcomes after injury. One plausible mechanism is that menopausal hormonal changes and HRT modulate neuroinflammatory or repair processes in ways that make the postmenopausal brain more sensitive to trauma. In this context, longer HRT exposure could exacerbate microstructural alterations following TBI, reflecting a potential synergistic interaction between exogenous hormones and injury-related processes. These findings underscore the importance of considering HRT timing and duration when evaluating neural outcomes after postmenopausal TBI.

Lastly, across our findings, a consistent pattern of anterior corpus callosum vulnerability emerged. TBI-related alterations were strongest in the genu, HRT effects in healthy women localized anteriorly, and the HRT duration interaction in postmenopausal TBI cases manifested specifically as increased genu MD. This convergence suggests that anterior regions, which contain late-myelinating fibers particularly susceptible to age-related decline (51), may represent sites where injury-related and hormone-related processes synergistically accelerate brain aging. Future studies with longitudinal designs, detailed HRT characterization, and multimodal imaging will be critical to disentangle these effects.

## Limitations

Several limitations warrant consideration. First, UK Biobank participants are healthier and more educated than the general population, with this selection bias especially evident among those who completed neuroimaging (52–54). This may attenuate effect estimates and reduce generalizability to more diverse populations or individuals with limited healthcare access. Second, our TBI ascertainment using ICD-10 hospital admission codes, while objective, has inherent limitations. Hospital records do not capture injury severity indicators such as Glasgow Coma Scale scores, duration of altered consciousness, or post-traumatic amnesia. Additionally, the administrative coding system records only initial diagnoses, preventing differentiation between single and recurrent injuries. Many individuals with TBI receive care exclusively in outpatient or primary care settings that are not included in hospital episode statistics, likely resulting in underrepresentation of less severe injuries in our sample. Third, the small pre-menopause TBI sample is a critical limitation affecting precision and generalizability of menopause timing comparisons and statistical power. Fourth, HRT data were self-reported and lacked specificity regarding preparation type, delivery method, dosing regimens, and temporal relationship to perimenopause and menopause onset. Fifth, the cross-sectional design with single-timepoint imaging prevents assessment of whether white matter changes represent static injury sequelae versus progressive degeneration, and whether trajectories differ by menopause timing or hormonal exposures. Sixth, our analysis focused exclusively on corpus callosum microstructure. Effects on other commissural tracts, association pathways, or subcortical gray matter regions remain unexamined. Similarly, we did not assess markers of cerebrovascular integrity or neuroinflammation that could provide mechanistic insights. Prospective studies with repeated neuroimaging, comprehensive injury characterization, detailed hormonal exposure documentation, and diverse populations are needed to clarify causal relationships and temporal dynamics.

## Conclusion

Females with a history of TBI exhibit persistent corpus callosum white matter alterations, with menopause timing and HRT exposure influencing these outcomes in complex, context-dependent ways. While HRT did not modify TBI effects in primary analyses, duration analyses suggest both potential benefits and risks, highlighting the importance of timing and cumulative exposure. Reproductive history influenced baseline white matter but did not confound TBI or HRT associations. These findings underscore the need for sex-specific, hormonally informed TBI research and longitudinal studies to clarify mechanisms and identify targeted interventions for women.

## Acknowledgments

This work was completed in part using the Discovery cluster, supported by Northeastern University’s Research Computing team.

## Author Contributions

### Consent for Participation

Written informed consent was obtained from all UK Biobank participants at the time of enrollment.

This study involved secondary analysis of de-identified data from UK Biobank conducted under application ID UKB project-GxVJqV0JfzGZQKFgg95b97ZP.

### Consent for Publication

Not applicable. This study uses de-identified data from UK Biobank, and no individual participant data are presented in this manuscript.

