## Supplementary Table 1 for "Traumatic Brain Injury Associated with Altered Corpus Callosum Microstructure in Females: Exploring the Roles of Menopause Timing and Hormone Therapy in UK Biobank"

**UK Biobank ICD-10 terms used to derive TBI status**

| | ICD-10 code |
|---|---|
| **Narrow-band traumatic brain injury (TBI)** | S020 |
| | S0200 |
| | S0201 |
| | S021 |
| | S0210 |
| | S0211 |
| | S027 |
| | S0270 |
| | S0271 |
| | S028 |
| | S0280 |
| | S0281 |
| | S029 |
| | S0290 |
| | S0291 |
| | S06 |
| | S060 |
| | S0600 |
| | S0601 |
| | S061 |
| | S0610 |
| | S0611 |
| | S062 |
| | S0620 |
| | S0621 |
| | S063 |
| | S0630 |
| | S0631 |
| | S064 |
| | S0640 |
| | S0641 |
| | S065 |
| | S0650 |
| | S0651 |
| | S066 |
| | S0660 |
| | S0661 |
| | S067 |
| | S0670 |

| | |
|---|---|
| | S0671 |
| | S068 |
| | S0680 |
| | S0681 |
| | S069 |
| | S0690 |
| | S0691 |
| | S07 |
| | S09 |
| | S097 |
| | S098 |
| | S099 |
| **Broad-band TBI (including narrow-band)** | G913 |
| | T90 |
| | T900 |
| | T901 |
| | T902 |
| | T905 |
| | T908 |
| | T909 |
| | S00 |
| | S000 |
| | S007 |
| | S008 |
| | S009 |
| | S01 |
| | S010 |
| | S011 |
| | S012 |
| | S013 |
| | S014 |
| | S015 |
| | S017 |
| | S018 |
| | S019 |
| | S02 |
| | S022 |
| | S0220 |
| | S0221 |
| | S023 |
| | S0230 |

S0231
S024
S0240
S0241
S026
S0260
S0261
S070
S071
S078
S079
S08
S080
S081
S088
S089
S090
S091
S092
V01
V010
V011
V019
V02
V020
V021
V029
V03
V030
V0308
V031
V0318
V0319
V039
V0399
V04
V040
V041
V049
V05
V050

V0509
V051
V059
V06
V060
V061
V069
V0691
V09
V090
V091
V092
V093
V099
V10
V100
V101
V102
V103
V104
V105
V109
V11
V110
V111
V112
V113
V114
V115
V119
V12
V120
V121
V122
V123
V124
V1248
V125
V129
V13
V130

V131
V132
V1328
V133
V134
V1349
V135
V139
V1399
V14
V140
V141
V142
V143
V144
V145
V149
V15
V150
V151
V152
V153
V154
V155
V159
V16
V160
V161
V162
V163
V164
V165
V169
V17
V170
V171
V172
V173
V174
V175
V179

V18
V180
V181
V182
V183
V184
V1849
V185
V189
V1899
V19
V190
V191
V192
V1929
V193
V194
V195
V196
V1969
V198
V199
V1999
V20
V200
V201
V202
V203
V204
V205
V209
V21
V210
V211
V212
V213
V214
V215
V219
V22
V220

V221
V222
V223
V224
V225
V229
V23
V230
V231
V232
V233
V234
V235
V239
V2399
V24
V240
V241
V242
V243
V244
V245
V249
V25
V250
V251
V252
V253
V254
V255
V259
V26
V260
V261
V262
V263
V264
V265
V269
V27
V270

V2709
V271
V272
V273
V274
V275
V279
V28
V280
V2809
V281
V282
V283
V284
V2849
V285
V289
V29
V290
V291
V292
V293
V294
V295
V296
V2969
V298
V2980
V299
V2999
V30
V300
V301
V302
V303
V304
V305
V306
V307
V309
V31

V310
V311
V312
V313
V314
V315
V316
V317
V319
V32
V320
V321
V322
V323
V324
V325
V326
V327
V329
V33
V330
V331
V332
V333
V334
V335
V336
V337
V339
V34
V340
V341
V342
V343
V344
V345
V346
V347
V349
V35
V350

V351
V352
V353
V354
V355
V356
V357
V359
V36
V360
V361
V362
V363
V364
V365
V366
V367
V369
V37
V370
V371
V372
V373
V374
V375
V376
V377
V379
V38
V380
V381
V382
V383
V384
V385
V386
V387
V389
V39
V390
V391

V392
V393
V394
V395
V396
V398
V399
V40
V400
V401
V402
V403
V404
V405
V406
V407
V409
V41
V410
V411
V412
V413
V414
V415
V416
V417
V419
V42
V420
V421
V422
V423
V424
V425
V4258
V426
V427
V429
V43
V430
V4309

V431
V432
V433
V434
V435
V4359
V436
V4369
V437
V439
V44
V440
V441
V442
V443
V444
V445
V446
V4469
V447
V449
V45
V450
V451
V452
V453
V454
V455
V456
V457
V459
V46
V460
V461
V462
V463
V464
V465
V466
V467
V469

V47
V470
V4709
V471
V472
V473
V474
V475
V4758
V4759
V476
V4769
V477
V479
V48
V480
V481
V482
V483
V484
V485
V4859
V486
V487
V489
V49
V490
V491
V492
V493
V494
V495
V496
V498
V4981
V499
V50
V500
V501
V502
V503

V504
V505
V506
V507
V509
V51
V510
V511
V512
V513
V514
V515
V516
V517
V519
V52
V520
V521
V522
V523
V524
V525
V526
V527
V529
V53
V530
V531
V532
V533
V534
V535
V536
V537
V539
V54
V540
V541
V542
V543
V544

V545
V546
V547
V549
V55
V550
V551
V552
V553
V554
V555
V556
V557
V559
V56
V560
V561
V562
V563
V564
V565
V566
V567
V569
V57
V570
V571
V572
V573
V574
V575
V576
V577
V579
V58
V580
V581
V582
V583
V584
V585

V586
V587
V589
V59
V590
V591
V592
V593
V594
V595
V596
V598
V599
V60
V600
V601
V602
V603
V604
V605
V606
V607
V609
V61
V610
V611
V612
V613
V614
V615
V616
V617
V619
V62
V620
V621
V622
V623
V624
V625
V626

V627
V629
V63
V630
V631
V632
V633
V634
V635
V636
V637
V639
V64
V640
V641
V642
V643
V644
V645
V646
V647
V649
V65
V650
V651
V652
V653
V654
V655
V656
V657
V659
V66
V660
V661
V662
V663
V664
V665
V666
V667

V669
V67
V670
V671
V672
V673
V674
V675
V676
V677
V679
V68
V680
V681
V682
V683
V684
V685
V686
V687
V689
V69
V690
V691
V692
V693
V694
V695
V696
V698
V699
V70
V700
V701
V702
V703
V704
V705
V706
V707
V709

V71
V710
V711
V712
V713
V714
V715
V716
V717
V719
V72
V720
V721
V722
V723
V724
V725
V726
V727
V729
V73
V730
V731
V732
V733
V734
V735
V736
V737
V739
V74
V740
V741
V742
V743
V744
V745
V746
V747
V749
V75

V750
V751
V752
V753
V754
V755
V756
V757
V759
V76
V760
V761
V762
V763
V764
V765
V766
V767
V769
V77
V770
V771
V772
V773
V774
V775
V776
V777
V779
V78
V780
V781
V782
V783
V784
V7849
V785
V786
V787
V789
V79

V790
V791
V792
V793
V794
V795
V796
V798
V7989
V799
V80
V800
V8000
V8001
V8009
V801
V802
V803
V804
V805
V806
V807
V808
V809
V81
V810
V811
V812
V813
V814
V815
V816
V817
V818
V819
V82
V820
V821
V822
V823
V824

V825
V826
V827
V828
V829
V83
V830
V831
V832
V833
V834
V835
V836
V837
V8372
V839
V84
V840
V841
V842
V843
V844
V845
V846
V847
V849
V85
V850
V851
V852
V853
V854
V8542
V855
V856
V857
V859
V86
V860
V861
V862

V863
V864
V865
V866
V867
V869
V87
V870
V871
V872
V873
V874
V875
V876
V877
V878
V879
V88
V880
V881
V882
V883
V884
V885
V886
V887
V888
V889
V89
V890
V891
V892
V893
V899
V90
V900
V901
V902
V903
V904
V905

V906
V907
V908
V909
V91
V910
V911
V912
V913
V914
V915
V916
V917
V918
V919
V92
V920
V921
V922
V923
V924
V925
V926
V927
V928
V929
V93
V930
V931
V932
V933
V9331
V934
V935
V936
V937
V938
V939
V94
V940
V941

V942
V943
V944
V945
V946
V947
V948
V949
V95
V950
V951
V952
V953
V954
V958
V959
V96
V960
V961
V962
V968
V969
V97
V970
V971
V972
V973
V978
V98
V99
W00
W000
W0008
W001
W002
W003
W004
W0042
W005
W006
W007

W008
W0089
W009
W0099
W01
W010
W0103
W0104
W0108
W0109
W011
W012
W0120
W0122
W0124
W0129
W013
W0130
W0131
W014
W0148
W0149
W015
W0153
W016
W017
W018
W0181
W0188
W0189
W019
W0190
W02
W020
W021
W022
W023
W0230
W0231
W024
W025

W026
W027
W028
W0281
W029
W03
W030
W031
W032
W033
W0330
W034
W035
W036
W037
W038
W0389
W039
W04
W040
W041
W042
W043
W044
W045
W046
W047
W048
W049
W05
W050
W051
W052
W053
W054
W055
W056
W057
W058
W059
W06

W060
W061
W062
W0628
W063
W064
W065
W066
W067
W068
W069
W07
W070
W071
W072
W0729
W073
W074
W075
W076
W077
W078
W079
W08
W080
W081
W082
W083
W084
W085
W086
W087
W088
W089
W09
W090
W091
W092
W093
W094
W095

W096

W097

W098

W099

W10

W100

W1003

W1004

W1008

W101

W102

W1029

W103

W104

W105

W1059

W106

W107

W108

W109

W1090

W1092

W11

W110

W1103

W1109

W111

W112

W113

W114

W115

W116

W117

W118

W1182

W119

W1199

W12

W120

W121

W122

W123
W124
W125
W126
W1262
W127
W128
W129
W1292
W1299
W13
W130
W1308
W131
W132
W133
W134
W135
W136
W137
W138
W1389
W139
W14
W140
W141
W142
W143
W144
W145
W146
W147
W148
W149
W15
W150
W151
W152
W153
W154
W155

W156
W157
W158
W159
W16
W160
W161
W162
W163
W164
W165
W166
W167
W168
W169
W17
W170
W1709
W171
W172
W173
W174
W175
W176
W177
W178
W1781
W179
W1799
W18
W180
W1804
W1809
W181
W182
W1822
W183
W184
W1848
W1849
W185

W186
W187
W188
W1889
W189
W1893
W1899
W19
W190
W1908
W1909
W191
W192
W193
W194
W195
W1959
W196
W197
W198
W1981
W199
W20
W200
W201
W202
W2022
W203
W204
W2049
W205
W2052
W206
W2062
W207
W208
W209
W2092
W2099
W21
W210

W211
W212
W213
W2130
W214
W215
W216
W217
W218
W219
W22
W220
W2203
W2209
W221
W222
W223
W224
W2241
W225
W226
W2262
W227
W228
W2288
W229
W23
W230
W2303
W2308
W2309
W231
W232
W233
W234
W235
W2352
W236
W2369
W237
W238

W2382
W239
W2392
W2399
W24
W240
W241
W242
W243
W244
W245
W246
W247
W248
W2482
W249
W25
W250
W2503
W2509
W251
W2518
W252
W253
W254
W255
W2552
W256
W2562
W257
W258
W259
W26
W260
W2603
W2608
W2609
W261
W262
W263
W264

W265
W2652
W266
W267
W268
W269
W2699
W27
W270
W2703
W2709
W271
W272
W273
W274
W275
W276
W277
W278
W279
W2792
W2799
W28
W280
W2800
W2801
W281
W282
W283
W284
W285
W286
W287
W288
W289
W29
W290
W2909
W291
W292
W293

W294
W295
W296
W2962
W297
W298
W2982
W299
W30
W300
W301
W302
W303
W304
W305
W306
W307
W308
W309
W31
W310
W3103
W311
W312
W313
W314
W315
W3152
W316
W3162
W317
W318
W3182
W3189
W319
W3192
W32
W320
W321
W322
W323

W324
W325
W326
W327
W328
W329
W33
W330
W331
W332
W333
W334
W335
W336
W337
W338
W339
W34
W340
W3409
W341
W342
W343
W344
W345
W346
W347
W348
W349
W35
W350
W351
W352
W353
W354
W355
W356
W357
W358
W359
W36

W360
W361
W362
W363
W364
W365
W366
W367
W368
W369
W37
W370
W371
W372
W373
W374
W375
W376
W377
W378
W379
W38
W380
W381
W382
W383
W384
W385
W386
W387
W388
W389
W39
W390
W391
W392
W393
W394
W395
W396
W397

W398
W399
W40
W400
W401
W402
W403
W404
W405
W406
W407
W408
W409
W41
W410
W411
W412
W413
W414
W415
W416
W417
W418
W419
W42
W420
W421
W422
W423
W424
W425
W426
W427
W428
W429
W43
W430
W431
W432
W433
W434

W435
W436
W437
W438
W439
W44
W440
W441
W442
W443
W444
W445
W4459
W446
W447
W448
W449
W4499
W45
W450
W4503
W4509
W451
W452
W453
W454
W455
W456
W457
W458
W459
W4599
W46
W460
W461
W462
W463
W464
W465
W466
W467

W468
W469
W49
W490
W491
W492
W493
W494
W495
W496
W497
W498
W499
W50
W500
W501
W502
W503
W5030
W504
W505
W506
W507
W508
W509
W51
W510
W5108
W511
W512
W513
W5130
W514
W515
W516
W517
W518
W5181
W519
W52
W520

W521
W522
W523
W524
W525
W526
W527
W528
W529
W53
W530
W531
W532
W533
W534
W535
W536
W537
W538
W539
W54
W540
W5408
W5409
W541
W542
W543
W544
W5449
W545
W546
W547
W548
W5481
W5488
W5489
W549
W5493
W5499
W55
W550

W551
W552
W553
W554
W555
W5558
W556
W557
W558
W559
W5599
W56
W560
W561
W562
W563
W564
W565
W566
W567
W568
W569
W57
W570
W571
W572
W573
W574
W575
W576
W577
W578
W579
W5799
W58
W580
W581
W582
W583
W584
W585

W586
W587
W588
W589
W59
W590
W591
W592
W593
W594
W595
W596
W597
W598
W599
W60
W600
W601
W602
W603
W604
W605
W606
W607
W608
W609
W64
W640
W641
W642
W643
W644
W645
W646
W647
W648
W649

**Meaning**

S02.0 Fracture of vault of skull

S02.00 Fracture of vault of skull (closed)

S02.01 Fracture of vault of skull (open)

S02.1 Fracture of base of skull

S02.10 Fracture of base of skull (closed)

S02.11 Fracture of base of skull (open)

S02.7 Multiple fractures involving skull and facial bones

S02.70 Multiple fractures involving skull and facial bones (closed)

S02.71 Multiple fractures involving skull and facial bones (open)

S02.8 Fractures of other skull and facial bones

S02.80 Fractures of other skull and facial bones (closed)

S02.81 Fractures of other skull and facial bones (open)

S02.9 Fracture of skull and facial bones, part unspecified

S02.90 Fracture of skull and facial bones, part unspecified (closed)

S02.91 Fracture of skull and facial bones, part unspecified (open)

S06 Intracranial injury

S06.0 Concussion

S06.00 Concussion (without open intracranial wound)

S06.01 Concussion (with open intracranial wound)

S06.1 Traumatic cerebral oedema

S06.10 Traumatic cerebral oedema (without open intracranial wound)

S06.11 Traumatic cerebral oedema (with open intracranial wound)

S06.2 Diffuse brain injury

S06.20 Diffuse brain injury (without open intracranial wound)

S06.21 Diffuse brain injury (with open intracranial wound)

S06.3 Focal brain injury

S06.30 Focal brain injury (without open intracranial wound)

S06.31 Focal brain injury (with open intracranial wound)

S06.4 Epidural haemorrhage

S06.40 Epidural haemorrhage (without open intracranial wound)

S06.41 Epidural haemorrhage (with open intracranial wound)

S06.5 Traumatic subdural haemorrhage

S06.50 Traumatic subdural haemorrhage (without open intracranial wound)

S06.51 Traumatic subdural haemorrhage (with open intracranial wound)

S06.6 Traumatic subarachnoid haemorrhage

S06.60 Traumatic subarachnoid haemorrhage (without open intracranial wound)

S06.61 Traumatic subarachnoid haemorrhage (with open intracranial wound)

S06.7 Intracranial injury with prolonged coma

S06.70 Intracranial injury with prolonged coma (without open intracranial wound)

S06.71 Intracranial injury with prolonged coma (with open intracranial wound)
S06.8 Other intracranial injuries
S06.80 Other intracranial injuries (without open intracranial wound)
S06.81 Other intracranial injuries (with open intracranial wound)
S06.9 Intracranial injury, unspecified
S06.90 Intracranial injury, unspecified (without open intracranial wound)
S06.91 Intracranial injury, unspecified (with open intracranial wound)
S07 Crushing injury of head
S09 Other and unspecified injuries of head
S09.7 Multiple injuries of head
S09.8 Other specified injuries of head
S09.9 Unspecified injury of head
G91.3 Posttraumatic hydrocephalus, unspecified
T90 Sequelae of injuries of head
T90.0 Sequelae of superficial injury of head
T90.1 Sequelae of open wound of head
T90.2 Sequelae of fracture of skull and facial bones
T90.5 Sequelae of intracranial injury
T90.8 Sequelae of other specified injuries of head
T90.9 Sequelae of unspecified injury of head
S00 Superficial injury of head
S00.0 Superficial injury of scalp
S00.7 Multiple superficial injuries of head
S00.8 Superficial injury of other parts of head
S00.9 Superficial injury of head, part unspecified
S01 Open wound of head
S01.0 Open wound of scalp
S01.1 Open wound of eyelid and periocular area
S01.2 Open wound of nose
S01.3 Open wound of ear
S01.4 Open wound of cheek and temporomandibular area
S01.5 Open wound of lip and oral cavity
S01.7 Multiple open wounds of head
S01.8 Open wound of other parts of head
S01.9 Open wound of head, part unspecified
S02 Fracture of skull and facial bones
S02.2 Fracture of nasal bones
S02.20 Fracture of nasal bones (closed)
S02.21 Fracture of nasal bones (open)
S02.3 Fracture of orbital floor
S02.30 Fracture of orbital floor (closed)

S02.31 Fracture of orbital floor (open)
S02.4 Fracture of malar and maxillary bones
S02.40 Fracture of malar and maxillary bones (closed)
S02.41 Fracture of malar and maxillary bones (open)
S02.6 Fracture of mandible
S02.60 Fracture of mandible (closed)
S02.61 Fracture of mandible (open)
S07.0 Crushing injury of face
S07.1 Crushing injury of skull
S07.8 Crushing injury of other parts of head
S07.9 Crushing injury of head, part unspecified
S08 Traumatic amputation of part of head
S08.0 Avulsion of scalp
S08.1 Traumatic amputation of ear
S08.8 Traumatic amputation of other parts of head
S08.9 Traumatic amputation of unspecified part of head
S09.0 Injury of blood vessels of head, not elsewhere classified
S09.1 Injury of muscle and tendon of head
S09.2 Traumatic rupture of ear drum
V01 Pedestrian injured in collision with pedal cycle
V01.0 Nontraffic accident
V01.1 Traffic accident
V01.9 Unspecified whether traffic or nontraffic accident
V02 Pedestrian injured in collision with two- or three-wheeled motor vehicle
V02.0 Nontraffic accident
V02.1 Traffic accident
V02.9 Unspecified whether traffic or nontraffic accident
V03 Pedestrian injured in collision with car, pick-up truck or van
V03.0 Nontraffic accident
V03.08 Pedestrian injured in collision with car, pick-up truck or van; nontraffic accident; while engaged in other
V03.1 Traffic accident
V03.18 Pedestrian injured in collision with car, pick-up truck or van; traffic accident; while engaged in other spe
V03.19 Pedestrian with other conveyance injured in collision with car, pick-up truck or van in traffic accident
V03.9 Unspecified whether traffic or nontraffic accident
V03.99 Pedestrian injured in collision with car, pick-up truck or van; unspecified whether traffic or nontraffic ac
V04 Pedestrian injured in collision with heavy transport vehicle or bus
V04.0 Nontraffic accident
V04.1 Traffic accident
V04.9 Unspecified whether traffic or nontraffic accident
V05 Pedestrian injured in collision with railway train or railway vehicle
V05.0 Nontraffic accident

V05.09 Pedestrian injured in collision with railway train or railway vehicle; nontraffic accident; during unspecifi
V05.1 Traffic accident
V05.9 Unspecified whether traffic or nontraffic accident
V06 Pedestrian injured in collision with other nonmotor vehicle
V06.0 Nontraffic accident
V06.1 Traffic accident
V06.9 Unspecified whether traffic or nontraffic accident
V06.91 Pedestrian injured in collision with other nonmotor vehicle: Unspecified whether traffic or nontraffic acc
V09 Pedestrian injured in other and unspecified transport accidents
V09.0 Pedestrian injured in nontraffic accident involving other and unspecified motor vehicles
V09.1 Pedestrian injured in unspecified nontraffic accident
V09.2 Pedestrian injured in traffic accident involving other and unspecified motor vehicles
V09.3 Pedestrian injured in unspecified traffic accident
V09.9 Pedestrian injured in unspecified transport accident
V10 Pedal cyclist injured in collision with pedestrian or animal
V10.0 Driver injured in nontraffic accident
V10.1 Passenger injured in nontraffic accident
V10.2 Unspecified pedal cyclist injured in nontraffic accident
V10.3 Person injured while boarding or alighting
V10.4 Driver injured in traffic accident
V10.5 Passenger injured in traffic accident
V10.9 Unspecified pedal cyclist injured in traffic accident
V11 Pedal cyclist injured in collision with other pedal cycle
V11.0 Driver injured in nontraffic accident
V11.1 Passenger injured in nontraffic accident
V11.2 Unspecified pedal cyclist injured in nontraffic accident
V11.3 Person injured while boarding or alighting
V11.4 Driver injured in traffic accident
V11.5 Passenger injured in traffic accident
V11.9 Unspecified pedal cyclist injured in traffic accident
V12 Pedal cyclist injured in collision with two- or three-wheeled motor vehicle
V12.0 Driver injured in nontraffic accident
V12.1 Passenger injured in nontraffic accident
V12.2 Unspecified pedal cyclist injured in nontraffic accident
V12.3 Person injured while boarding or alighting
V12.4 Driver injured in traffic accident
V12.48 Pedal cyclist injured in collision with two- or three-wheeled motor vehicle; driver injured in traffic accid
V12.5 Passenger injured in traffic accident
V12.9 Unspecified pedal cyclist injured in traffic accident
V13 Pedal cyclist injured in collision with car, pick-up truck or van
V13.0 Driver injured in nontraffic accident

V13.1 Passenger injured in nontraffic accident
V13.2 Unspecified pedal cyclist injured in nontraffic accident
V13.28 Pedal cyclist injured in collision with car, pick-up truck or van; unspecified pedal cyclist injured in nontr
V13.3 Person injured while boarding or alighting
V13.4 Driver injured in traffic accident
V13.49 Pedal cyclist injured in collision with car, pick-up truck or van; driver injured in traffic accident ; during
V13.5 Passenger injured in traffic accident
V13.9 Unspecified pedal cyclist injured in traffic accident
V13.99 Pedal cyclist injured in collision with car, pick-up truck or van; unspecified pedal cyclist injured in traffi
V14 Pedal cyclist injured in collision with heavy transport vehicle or bus
V14.0 Driver injured in nontraffic accident
V14.1 Passenger injured in nontraffic accident
V14.2 Unspecified pedal cyclist injured in nontraffic accident
V14.3 Person injured while boarding or alighting
V14.4 Driver injured in traffic accident
V14.5 Passenger injured in traffic accident
V14.9 Unspecified pedal cyclist injured in traffic accident
V15 Pedal cyclist injured in collision with railway train or railway vehicle
V15.0 Driver injured in nontraffic accident
V15.1 Passenger injured in nontraffic accident
V15.2 Unspecified pedal cyclist injured in nontraffic accident
V15.3 Person injured while boarding or alighting
V15.4 Driver injured in traffic accident
V15.5 Passenger injured in traffic accident
V15.9 Unspecified pedal cyclist injured in traffic accident
V16 Pedal cyclist injured in collision with other nonmotor vehicle
V16.0 Driver injured in nontraffic accident
V16.1 Passenger injured in nontraffic accident
V16.2 Unspecified pedal cyclist injured in nontraffic accident
V16.3 Person injured while boarding or alighting
V16.4 Driver injured in traffic accident
V16.5 Passenger injured in traffic accident
V16.9 Unspecified pedal cyclist injured in traffic accident
V17 Pedal cyclist injured in collision with fixed or stationary object
V17.0 Driver injured in nontraffic accident
V17.1 Passenger injured in nontraffic accident
V17.2 Unspecified pedal cyclist injured in nontraffic accident
V17.3 Person injured while boarding or alighting
V17.4 Driver injured in traffic accident
V17.5 Passenger injured in traffic accident
V17.9 Unspecified pedal cyclist injured in traffic accident

V18 Pedal cyclist injured in noncollision transport accident
V18.0 Driver injured in nontraffic accident
V18.1 Passenger injured in nontraffic accident
V18.2 Unspecified pedal cyclist injured in nontraffic accident
V18.3 Person injured while boarding or alighting
V18.4 Driver injured in traffic accident
V18.49 Pedal cyclist injured in noncollision transport accident; driver injured in traffic accident ; during unspeci
V18.5 Passenger injured in traffic accident
V18.9 Unspecified pedal cyclist injured in traffic accident
V18.99 Pedal cyclist injured in noncollision transport accident; unspecified pedal cyclist injured in traffic accide
V19 Pedal cyclist injured in other and unspecified transport accidents
V19.0 Driver injured in collision with other and unspecified motor vehicles in nontraffic accident
V19.1 Passenger injured in collision with other and unspecified motor vehicles in nontraffic accident
V19.2 Unspecified pedal cyclist injured in collision with other and unspecified motor vehicles in nontraffic accic
V19.29 Unspecified pedal cyclist injured in collision with other and unspecified motor vehicles in nontraffic acc
V19.3 Pedal cyclist [any] injured in an unspecified nontraffic accident
V19.4 Driver injured in collision with other and unspecified motor vehicles in traffic accident
V19.5 Passenger injured in collision with other and unspecified motor vehicles in traffic accident
V19.6 Unspecified pedal cyclist injured in collision with other and unspecified motor vehicles in traffic accident
V19.69 Unspecified pedal cyclist injured in collision with other and unspecified motor vehicles in traffic accider
V19.8 Pedal cyclist [any] injured in other specified transport accidents
V19.9 Pedal cyclist [any] injured in unspecified traffic accident
V19.99 Pedal cyclist [any] injured in unspecified traffic accident; during unspecified activity
V20 Motorcycle rider injured in collision with pedestrian or animal
V20.0 Driver injured in nontraffic accident
V20.1 Passenger injured in nontraffic accident
V20.2 Unspecified motorcycle rider injured in nontraffic accident
V20.3 Person injured while boarding or alighting
V20.4 Driver injured in traffic accident
V20.5 Passenger injured in traffic accident
V20.9 Unspecified motorcycle rider injured in traffic accident
V21 Motorcycle rider injured in collision with pedal cycle
V21.0 Driver injured in nontraffic accident
V21.1 Passenger injured in nontraffic accident
V21.2 Unspecified motorcycle rider injured in nontraffic accident
V21.3 Person injured while boarding or alighting
V21.4 Driver injured in traffic accident
V21.5 Passenger injured in traffic accident
V21.9 Unspecified motorcycle rider injured in traffic accident
V22 Motorcycle rider injured in collision with two- or three-wheeled motor vehicle
V22.0 Driver injured in nontraffic accident

V22.1 Passenger injured in nontraffic accident
V22.2 Unspecified motorcycle rider injured in nontraffic accident
V22.3 Person injured while boarding or alighting
V22.4 Driver injured in traffic accident
V22.5 Passenger injured in traffic accident
V22.9 Unspecified motorcycle rider injured in traffic accident
V23 Motorcycle rider injured in collision with car, pick-up truck or van
V23.0 Driver injured in nontraffic accident
V23.1 Passenger injured in nontraffic accident
V23.2 Unspecified motorcycle rider injured in nontraffic accident
V23.3 Person injured while boarding or alighting
V23.4 Driver injured in traffic accident
V23.5 Passenger injured in traffic accident
V23.9 Unspecified motorcycle rider injured in traffic accident
V23.99 Motorcycle rider injured in collision with car, pick-up truck or van, Unspecified motorcycle rider injurec
V24 Motorcycle rider injured in collision with heavy transport vehicle or bus
V24.0 Driver injured in nontraffic accident
V24.1 Passenger injured in nontraffic accident
V24.2 Unspecified motorcycle rider injured in nontraffic accident
V24.3 Person injured while boarding or alighting
V24.4 Driver injured in traffic accident
V24.5 Passenger injured in traffic accident
V24.9 Unspecified motorcycle rider injured in traffic accident
V25 Motorcycle rider injured in collision with railway train or railway vehicle
V25.0 Driver injured in nontraffic accident
V25.1 Passenger injured in nontraffic accident
V25.2 Unspecified motorcycle rider injured in nontraffic accident
V25.3 Person injured while boarding or alighting
V25.4 Driver injured in traffic accident
V25.5 Passenger injured in traffic accident
V25.9 Unspecified motorcycle rider injured in traffic accident
V26 Motorcycle rider injured in collision with other nonmotor vehicle
V26.0 Driver injured in nontraffic accident
V26.1 Passenger injured in nontraffic accident
V26.2 Unspecified motorcycle rider injured in nontraffic accident
V26.3 Person injured while boarding or alighting
V26.4 Driver injured in traffic accident
V26.5 Passenger injured in traffic accident
V26.9 Unspecified motorcycle rider injured in traffic accident
V27 Motorcycle rider injured in collision with fixed or stationary object
V27.0 Driver injured in nontraffic accident

V27.09 Motorcycle rider injured in collision with fixed or stationary object; driver injured in nontraffic accident
V27.1 Passenger injured in nontraffic accident
V27.2 Unspecified motorcycle rider injured in nontraffic accident
V27.3 Person injured while boarding or alighting
V27.4 Driver injured in traffic accident
V27.5 Passenger injured in traffic accident
V27.9 Unspecified motorcycle rider injured in traffic accident
V28 Motorcycle rider injured in noncollision transport accident
V28.0 Driver injured in nontraffic accident
V28.09 Motorcycle rider injured in noncollision transport accident; driver injured in nontraffic accident; during [illegible]
V28.1 Passenger injured in nontraffic accident
V28.2 Unspecified motorcycle rider injured in nontraffic accident
V28.3 Person injured while boarding or alighting
V28.4 Driver injured in traffic accident
V28.49 Motorcycle rider injured in noncollision transport accident; driver injured in traffic accident; during uns[illegible]
V28.5 Passenger injured in traffic accident
V28.9 Unspecified motorcycle rider injured in traffic accident
V29 Motorcycle rider injured in other and unspecified transport accidents
V29.0 Driver injured in collision with other and unspecified motor vehicles in nontraffic accident
V29.1 Passenger injured in collision with other and unspecified motor vehicles in nontraffic accident
V29.2 Unspecified motorcycle rider injured in collision with other and unspecified motor vehicles in nontraffic [illegible]
V29.3 Motorcycle rider [any] injured in unspecified nontraffic accident
V29.4 Driver injured in collision with other and unspecified motor vehicles in traffic accident
V29.5 Passenger injured in collision with other and unspecified motor vehicles in traffic accident
V29.6 Unspecified motorcycle rider injured in collision with other and unspecified motor vehicles in traffic acci[illegible]
V29.69 Unspecified motorcycle rider injured in collision with other and unspecified motor vehicles in traffic acc[illegible]
V29.8 Motorcycle rider [any] injured in other specified transport accidents
V29.80 Motorcycle rider [any] injured in other specified transport accidents; while engaged in sports activity
V29.9 Motorcycle rider [any] injured in unspecified traffic accident
V29.99 Motorcycle rider [any] injured in unspecified traffic accident; during unspecified activity
V30 Occupant of three-wheeled motor vehicle injured in collision with pedestrian or animal
V30.0 Driver injured in nontraffic accident
V30.1 Passenger injured in nontraffic accident
V30.2 Person on outside of vehicle injured in nontraffic accident
V30.3 Unspecified occupant of three-wheeled motor vehicle injured in nontraffic accident
V30.4 Person injured while boarding or alighting
V30.5 Driver injured in traffic accident
V30.6 Passenger injured in traffic accident
V30.7 Person on outside of vehicle injured in traffic accident
V30.9 Unspecified occupant of three-wheeled motor vehicle injured in traffic accident
V31 Occupant of three-wheeled motor vehicle injured in collision with pedal cycle

V31.0 Driver injured in nontraffic accident
V31.1 Passenger injured in nontraffic accident
V31.2 Person on outside of vehicle injured in nontraffic accident
V31.3 Unspecified occupant of three-wheeled motor vehicle injured in nontraffic accident
V31.4 Person injured while boarding or alighting
V31.5 Driver injured in traffic accident
V31.6 Passenger injured in traffic accident
V31.7 Person on outside of vehicle injured in traffic accident
V31.9 Unspecified occupant of three-wheeled motor vehicle injured in traffic accident
V32 Occupant of three-wheeled motor vehicle injured in collision with two- or three-wheeled motor vehicle
V32.0 Driver injured in nontraffic accident
V32.1 Passenger injured in nontraffic accident
V32.2 Person on outside of vehicle injured in nontraffic accident
V32.3 Unspecified occupant of three-wheeled motor vehicle injured in nontraffic accident
V32.4 Person injured while boarding or alighting
V32.5 Driver injured in traffic accident
V32.6 Passenger injured in traffic accident
V32.7 Person on outside of vehicle injured in traffic accident
V32.9 Unspecified occupant of three-wheeled motor vehicle injured in traffic accident
V33 Occupant of three-wheeled motor vehicle injured in collision with car, pick-up truck or van
V33.0 Driver injured in nontraffic accident
V33.1 Passenger injured in nontraffic accident
V33.2 Person on outside of vehicle injured in nontraffic accident
V33.3 Unspecified occupant of three-wheeled motor vehicle injured in nontraffic accident
V33.4 Person injured while boarding or alighting
V33.5 Driver injured in traffic accident
V33.6 Passenger injured in traffic accident
V33.7 Person on outside of vehicle injured in traffic accident
V33.9 Unspecified occupant of three-wheeled motor vehicle injured in traffic accident
V34 Occupant of three-wheeled motor vehicle injured in collision with heavy transport vehicle or bus
V34.0 Driver injured in nontraffic accident
V34.1 Passenger injured in nontraffic accident
V34.2 Person on outside of vehicle injured in nontraffic accident
V34.3 Unspecified occupant of three-wheeled motor vehicle injured in nontraffic accident
V34.4 Person injured while boarding or alighting
V34.5 Driver injured in traffic accident
V34.6 Passenger injured in traffic accident
V34.7 Person on outside of vehicle injured in traffic accident
V34.9 Unspecified occupant of three-wheeled motor vehicle injured in traffic accident
V35 Occupant of three-wheeled motor vehicle injured in collision with railway train or railway vehicle
V35.0 Driver injured in nontraffic accident

V35.1 Passenger injured in nontraffic accident
V35.2 Person on outside of vehicle injured in nontraffic accident
V35.3 Unspecified occupant of three-wheeled motor vehicle injured in nontraffic accident
V35.4 Person injured while boarding or alighting
V35.5 Driver injured in traffic accident
V35.6 Passenger injured in traffic accident
V35.7 Person on outside of vehicle injured in traffic accident
V35.9 Unspecified occupant of three-wheeled motor vehicle injured in traffic accident
V36 Occupant of three-wheeled motor vehicle injured in collision with other nonmotor vehicle
V36.0 Driver injured in nontraffic accident
V36.1 Passenger injured in nontraffic accident
V36.2 Person on outside of vehicle injured in nontraffic accident
V36.3 Unspecified occupant of three-wheeled motor vehicle injured in nontraffic accident
V36.4 Person injured while boarding or alighting
V36.5 Driver injured in traffic accident
V36.6 Passenger injured in traffic accident
V36.7 Person on outside of vehicle injured in traffic accident
V36.9 Unspecified occupant of three-wheeled motor vehicle injured in traffic accident
V37 Occupant of three-wheeled motor vehicle injured in collision with fixed or stationary object
V37.0 Driver injured in nontraffic accident
V37.1 Passenger injured in nontraffic accident
V37.2 Person on outside of vehicle injured in nontraffic accident
V37.3 Unspecified occupant of three-wheeled motor vehicle injured in nontraffic accident
V37.4 Person injured while boarding or alighting
V37.5 Driver injured in traffic accident
V37.6 Passenger injured in traffic accident
V37.7 Person on outside of vehicle injured in traffic accident
V37.9 Unspecified occupant of three-wheeled motor vehicle injured in traffic accident
V38 Occupant of three-wheeled motor vehicle injured in noncollision transport accident
V38.0 Driver injured in nontraffic accident
V38.1 Passenger injured in nontraffic accident
V38.2 Person on outside of vehicle injured in nontraffic accident
V38.3 Unspecified occupant of three-wheeled motor vehicle injured in nontraffic accident
V38.4 Person injured while boarding or alighting
V38.5 Driver injured in traffic accident
V38.6 Passenger injured in traffic accident
V38.7 Person on outside of vehicle injured in traffic accident
V38.9 Unspecified occupant of three-wheeled motor vehicle injured in traffic accident
V39 Occupant of three-wheeled motor vehicle injured in other and unspecified transport accidents
V39.0 Driver injured in collision with other and unspecified motor vehicles in nontraffic accident
V39.1 Passenger injured in collision with other and unspecified motor vehicles in nontraffic accident

V39.2 Unspecified occupant of three-wheeled motor vehicle injured in collision with other and unspecified moto
V39.3 Occupant [any] of three-wheeled motor vehicle injured in unspecified nontraffic accident
V39.4 Driver injured in collision with other and unspecified motor vehicles in traffic accident
V39.5 Passenger injured in collision with other and unspecified motor vehicles in traffic accident
V39.6 Unspecified occupant of three-wheeled motor vehicle injured in collision with other and unspecified moto
V39.8 Occupant [any] of three-wheeled motor vehicle injured in other specified transport accidents
V39.9 Occupant [any] of three-wheeled motor vehicle injured in unspecified traffic accident
V40 Car occupant injured in collision with pedestrian or animal
V40.0 Driver injured in nontraffic accident
V40.1 Passenger injured in nontraffic accident
V40.2 Person on outside of vehicle injured in nontraffic accident
V40.3 Unspecified car occupant injured in nontraffic accident
V40.4 Person injured while boarding or alighting
V40.5 Driver injured in traffic accident
V40.6 Passenger injured in traffic accident
V40.7 Person on outside of vehicle injured in traffic accident
V40.9 Unspecified car occupant injured in traffic accident
V41 Car occupant injured in collision with pedal cycle
V41.0 Driver injured in nontraffic accident
V41.1 Passenger injured in nontraffic accident
V41.2 Person on outside of vehicle injured in nontraffic accident
V41.3 Unspecified car occupant injured in nontraffic accident
V41.4 Person injured while boarding or alighting
V41.5 Driver injured in traffic accident
V41.6 Passenger injured in traffic accident
V41.7 Person on outside of vehicle injured in traffic accident
V41.9 Unspecified car occupant injured in traffic accident
V42 Car occupant injured in collision with two- or three-wheeled motor vehicle
V42.0 Driver injured in nontraffic accident
V42.1 Passenger injured in nontraffic accident
V42.2 Person on outside of vehicle injured in nontraffic accident
V42.3 Unspecified car occupant injured in nontraffic accident
V42.4 Person injured while boarding or alighting
V42.5 Driver injured in traffic accident
V42.58 Car occupant injured in collision with two- or three-wheeled motor vehicle; driver injured in traffic accic
V42.6 Passenger injured in traffic accident
V42.7 Person on outside of vehicle injured in traffic accident
V42.9 Unspecified car occupant injured in traffic accident
V43 Car occupant injured in collision with car, pick-up truck or van
V43.0 Driver injured in nontraffic accident
V43.09 Car occupant injured in collision with car, pick-up truck or van; driver injured in nontraffic accident; dur

V43.1 Passenger injured in nontraffic accident
V43.2 Person on outside of vehicle injured in nontraffic accident
V43.3 Unspecified car occupant injured in nontraffic accident
V43.4 Person injured while boarding or alighting
V43.5 Driver injured in traffic accident
V43.59 Car occupant injured in collision with car, pick-up truck or van; driver injured in traffic accident; during
V43.6 Passenger injured in traffic accident
V43.69 Car occupant injured in collision with car, pick-up truck or van; passenger injured in traffic accident; du
V43.7 Person on outside of vehicle injured in traffic accident
V43.9 Unspecified car occupant injured in traffic accident
V44 Car occupant injured in collision with heavy transport vehicle or bus
V44.0 Driver injured in nontraffic accident
V44.1 Passenger injured in nontraffic accident
V44.2 Person on outside of vehicle injured in nontraffic accident
V44.3 Unspecified car occupant injured in nontraffic accident
V44.4 Person injured while boarding or alighting
V44.5 Driver injured in traffic accident
V44.6 Passenger injured in traffic accident
V44.69 Car occupant injured in collision with heavy transport vehicle or bus; passenger injured in traffic accide
V44.7 Person on outside of vehicle injured in traffic accident
V44.9 Unspecified car occupant injured in traffic accident
V45 Car occupant injured in collision with railway train or railway vehicle
V45.0 Driver injured in nontraffic accident
V45.1 Passenger injured in nontraffic accident
V45.2 Person on outside of vehicle injured in nontraffic accident
V45.3 Unspecified car occupant injured in nontraffic accident
V45.4 Person injured while boarding or alighting
V45.5 Driver injured in traffic accident
V45.6 Passenger injured in traffic accident
V45.7 Person on outside of vehicle injured in traffic accident
V45.9 Unspecified car occupant injured in traffic accident
V46 Car occupant injured in collision with other nonmotor vehicle
V46.0 Driver injured in nontraffic accident
V46.1 Passenger injured in nontraffic accident
V46.2 Person on outside of vehicle injured in nontraffic accident
V46.3 Unspecified car occupant injured in nontraffic accident
V46.4 Person injured while boarding or alighting
V46.5 Driver injured in traffic accident
V46.6 Passenger injured in traffic accident
V46.7 Person on outside of vehicle injured in traffic accident
V46.9 Unspecified car occupant injured in traffic accident

V47 Car occupant injured in collision with fixed or stationary object
V47.0 Driver injured in nontraffic accident
V47.09 Car occupant injured in collision with fixed or stationary object; driver injured in nontraffic accident; du
V47.1 Passenger injured in nontraffic accident
V47.2 Person on outside of vehicle injured in nontraffic accident
V47.3 Unspecified car occupant injured in nontraffic accident
V47.4 Person injured while boarding or alighting
V47.5 Driver injured in traffic accident
V47.58 Car occupant injured in collision with fixed or stationary object; driver injured in traffic accident; while
V47.59 Car occupant injured in collision with fixed or stationary object; driver injured in traffic accident; during
V47.6 Passenger injured in traffic accident
V47.69 Car occupant injured in collision with fixed or stationary object; passenger injured in traffic accident; du
V47.7 Person on outside of vehicle injured in traffic accident
V47.9 Unspecified car occupant injured in traffic accident
V48 Car occupant injured in noncollision transport accident
V48.0 Driver injured in nontraffic accident
V48.1 Passenger injured in nontraffic accident
V48.2 Person on outside of vehicle injured in nontraffic accident
V48.3 Unspecified car occupant injured in nontraffic accident
V48.4 Person injured while boarding or alighting
V48.5 Driver injured in traffic accident
V48.59 Car occupant injured in noncollision transport accident; driver injured in traffic accident; during unspeci
V48.6 Passenger injured in traffic accident
V48.7 Person on outside of vehicle injured in traffic accident
V48.9 Unspecified car occupant injured in traffic accident
V49 Car occupant injured in other and unspecified transport accidents
V49.0 Driver injured in collision with other and unspecified motor vehicles in nontraffic accident
V49.1 Passenger injured in collision with other and unspecified motor vehicles in nontraffic accident
V49.2 Unspecified car occupant injured in collision with other and unspecified motor vehicles in nontraffic acci
V49.3 Car occupant [any] injured in unspecified nontraffic accident
V49.4 Driver injured in collision with other and unspecified motor vehicles in traffic accident
V49.5 Passenger injured in collision with other and unspecified motor vehicles in traffic accident
V49.6 Unspecified car occupant injured in collision with other and unspecified motor vehicles in traffic accident
V49.8 Car occupant [any] injured in other specified transport accidents
V49.81 Car occupant injured in trnsp accident w military vehicle (Car occupant (driver) (passenger) injured in tr
V49.9 Car occupant [any] injured in unspecified traffic accident
V50 Occupant of pick-up truck or van injured in collision with pedestrian or animal
V50.0 Driver injured in nontraffic accident
V50.1 Passenger injured in nontraffic accident
V50.2 Person on outside of vehicle injured in nontraffic accident
V50.3 Unspecified occupant of pick-up truck or van injured in nontraffic accident

V50.4 Person injured while boarding or alighting
V50.5 Driver injured in traffic accident
V50.6 Passenger injured in traffic accident
V50.7 Person on outside of vehicle injured in traffic accident
V50.9 Unspecified occupant of pick-up truck or van injured in traffic accident
V51 Occupant of pick-up truck or van injured in collision with pedal cycle
V51.0 Driver injured in nontraffic accident
V51.1 Passenger injured in nontraffic accident
V51.2 Person on outside of vehicle injured in nontraffic accident
V51.3 Unspecified occupant of pick-up truck or van injured in nontraffic accident
V51.4 Person injured while boarding or alighting
V51.5 Driver injured in traffic accident
V51.6 Passenger injured in traffic accident
V51.7 Person on outside of vehicle injured in traffic accident
V51.9 Unspecified occupant of pick-up truck or van injured in traffic accident
V52 Occupant of pick-up truck or van injured in collision with two- or three-wheeled motor vehicle
V52.0 Driver injured in nontraffic accident
V52.1 Passenger injured in nontraffic accident
V52.2 Person on outside of vehicle injured in nontraffic accident
V52.3 Unspecified occupant of pick-up truck or van injured in nontraffic accident
V52.4 Person injured while boarding or alighting
V52.5 Driver injured in traffic accident
V52.6 Passenger injured in traffic accident
V52.7 Person on outside of vehicle injured in traffic accident
V52.9 Unspecified occupant of pick-up truck or van injured in traffic accident
V53 Occupant of pick-up truck or van injured in collision with car, pick-up truck or van
V53.0 Driver injured in nontraffic accident
V53.1 Passenger injured in nontraffic accident
V53.2 Person on outside of vehicle injured in nontraffic accident
V53.3 Unspecified occupant of pick-up truck or van injured in nontraffic accident
V53.4 Person injured while boarding or alighting
V53.5 Driver injured in traffic accident
V53.6 Passenger injured in traffic accident
V53.7 Person on outside of vehicle injured in traffic accident
V53.9 Unspecified occupant of pick-up truck or van injured in traffic accident
V54 Occupant of pick-up truck or van injured in collision with heavy transport vehicle or bus
V54.0 Driver injured in nontraffic accident
V54.1 Passenger injured in nontraffic accident
V54.2 Person on outside of vehicle injured in nontraffic accident
V54.3 Unspecified occupant of pick-up truck or van injured in nontraffic accident
V54.4 Person injured while boarding or alighting

V54.5 Driver injured in traffic accident
V54.6 Passenger injured in traffic accident
V54.7 Person on outside of vehicle injured in traffic accident
V54.9 Unspecified occupant of pick-up truck or van injured in traffic accident
V55 Occupant of pick-up truck or van injured in collision with railway train or railway vehicle
V55.0 Driver injured in nontraffic accident
V55.1 Passenger injured in nontraffic accident
V55.2 Person on outside of vehicle injured in nontraffic accident
V55.3 Unspecified occupant of pick-up truck or van injured in nontraffic accident
V55.4 Person injured while boarding or alighting
V55.5 Driver injured in traffic accident
V55.6 Passenger injured in traffic accident
V55.7 Person on outside of vehicle injured in traffic accident
V55.9 Unspecified occupant of pick-up truck or van injured in traffic accident
V56 Occupant of pick-up truck or van injured in collision with other nonmotor vehicle
V56.0 Driver injured in nontraffic accident
V56.1 Passenger injured in nontraffic accident
V56.2 Person on outside of vehicle injured in nontraffic accident
V56.3 Unspecified occupant of pick-up truck or van injured in nontraffic accident
V56.4 Person injured while boarding or alighting
V56.5 Driver injured in traffic accident
V56.6 Passenger injured in traffic accident
V56.7 Person on outside of vehicle injured in traffic accident
V56.9 Unspecified occupant of pick-up truck or van injured in traffic accident
V57 Occupant of pick-up truck or van injured in collision with fixed or stationary object
V57.0 Driver injured in nontraffic accident
V57.1 Passenger injured in nontraffic accident
V57.2 Person on outside of vehicle injured in nontraffic accident
V57.3 Unspecified occupant of pick-up truck or van injured in nontraffic accident
V57.4 Person injured while boarding or alighting
V57.5 Driver injured in traffic accident
V57.6 Passenger injured in traffic accident
V57.7 Person on outside of vehicle injured in traffic accident
V57.9 Unspecified occupant of pick-up truck or van injured in traffic accident
V58 Occupant of pick-up truck or van injured in noncollision transport accident
V58.0 Driver injured in nontraffic accident
V58.1 Passenger injured in nontraffic accident
V58.2 Person on outside of vehicle injured in nontraffic accident
V58.3 Unspecified occupant of pick-up truck or van injured in nontraffic accident
V58.4 Person injured while boarding or alighting
V58.5 Driver injured in traffic accident

V58.6 Passenger injured in traffic accident
V58.7 Person on outside of vehicle injured in traffic accident
V58.9 Unspecified occupant of pick-up truck or van injured in traffic accident
V59 Occupant of pick-up truck or van injured in other and unspecified transport accidents
V59.0 Driver injured in collision with other and unspecified motor vehicles in nontraffic accident
V59.1 Passenger injured in collision with other and unspecified motor vehicles in nontraffic accident
V59.2 Unspecified occupant of pick-up truck or van injured in collision with other and unspecified motor vehicl[illegible]
V59.3 Occupant [any] of pick-up truck or van injured in unspecified nontraffic accident
V59.4 Driver injured in collision with other and unspecified motor vehicles in traffic accident
V59.5 Passenger injured in collision with other and unspecified motor vehicles in traffic accident
V59.6 Unspecified occupant of pick-up truck or van injured in collision with other and unspecified motor vehicl[illegible]
V59.8 Occupant [any] of pick-up truck or van injured in other specified transport accidents
V59.9 Occupant [any] of pick-up truck or van injured in unspecified traffic accident
V60 Occupant of heavy transport vehicle injured in collision with pedestrian or animal
V60.0 Driver injured in nontraffic accident
V60.1 Passenger injured in nontraffic accident
V60.2 Person on outside of vehicle injured in nontraffic accident
V60.3 Unspecified occupant of heavy transport vehicle injured in nontraffic accident
V60.4 Person injured while boarding or alighting
V60.5 Driver injured in traffic accident
V60.6 Passenger injured in traffic accident
V60.7 Person on outside of vehicle injured in traffic accident
V60.9 Unspecified occupant of heavy transport vehicle injured in traffic accident
V61 Occupant of heavy transport vehicle injured in collision with pedal cycle
V61.0 Driver injured in nontraffic accident
V61.1 Passenger injured in nontraffic accident
V61.2 Person on outside of vehicle injured in nontraffic accident
V61.3 Unspecified occupant of heavy transport vehicle injured in nontraffic accident
V61.4 Person injured while boarding or alighting
V61.5 Driver injured in traffic accident
V61.6 Passenger injured in traffic accident
V61.7 Person on outside of vehicle injured in traffic accident
V61.9 Unspecified occupant of heavy transport vehicle injured in traffic accident
V62 Occupant of heavy transport vehicle injured in collision with two- or three-wheeled motor vehicle
V62.0 Driver injured in nontraffic accident
V62.1 Passenger injured in nontraffic accident
V62.2 Person on outside of vehicle injured in nontraffic accident
V62.3 Unspecified occupant of heavy transport vehicle injured in nontraffic accident
V62.4 Person injured while boarding or alighting
V62.5 Driver injured in traffic accident
V62.6 Passenger injured in traffic accident

V62.7 Person on outside of vehicle injured in traffic accident
V62.9 Unspecified occupant of heavy transport vehicle injured in traffic accident
V63 Occupant of heavy transport vehicle injured in collision with car, pick-up truck or van
V63.0 Driver injured in nontraffic accident
V63.1 Passenger injured in nontraffic accident
V63.2 Person on outside of vehicle injured in nontraffic accident
V63.3 Unspecified occupant of heavy transport vehicle injured in nontraffic accident
V63.4 Person injured while boarding or alighting
V63.5 Driver injured in traffic accident
V63.6 Passenger injured in traffic accident
V63.7 Person on outside of vehicle injured in traffic accident
V63.9 Unspecified occupant of heavy transport vehicle injured in traffic accident
V64 Occupant of heavy transport vehicle injured in collision with heavy transport vehicle or bus
V64.0 Driver injured in nontraffic accident
V64.1 Passenger injured in nontraffic accident
V64.2 Person on outside of vehicle injured in nontraffic accident
V64.3 Unspecified occupant of heavy transport vehicle injured in nontraffic accident
V64.4 Person injured while boarding or alighting
V64.5 Driver injured in traffic accident
V64.6 Passenger injured in traffic accident
V64.7 Person on outside of vehicle injured in traffic accident
V64.9 Unspecified occupant of heavy transport vehicle injured in traffic accident
V65 Occupant of heavy transport vehicle injured in collision with railway train or railway vehicle
V65.0 Driver injured in nontraffic accident
V65.1 Passenger injured in nontraffic accident
V65.2 Person on outside of vehicle injured in nontraffic accident
V65.3 Unspecified occupant of heavy transport vehicle injured in nontraffic accident
V65.4 Person injured while boarding or alighting
V65.5 Driver injured in traffic accident
V65.6 Passenger injured in traffic accident
V65.7 Person on outside of vehicle injured in traffic accident
V65.9 Unspecified occupant of heavy transport vehicle injured in traffic accident
V66 Occupant of heavy transport vehicle injured in collision with other nonmotor vehicle
V66.0 Driver injured in nontraffic accident
V66.1 Passenger injured in nontraffic accident
V66.2 Person on outside of vehicle injured in nontraffic accident
V66.3 Unspecified occupant of heavy transport vehicle injured in nontraffic accident
V66.4 Person injured while boarding or alighting
V66.5 Driver injured in traffic accident
V66.6 Passenger injured in traffic accident
V66.7 Person on outside of vehicle injured in traffic accident

V66.9 Unspecified occupant of heavy transport vehicle injured in traffic accident
V67 Occupant of heavy transport vehicle injured in collision with fixed or stationary object
V67.0 Driver injured in nontraffic accident
V67.1 Passenger injured in nontraffic accident
V67.2 Person on outside of vehicle injured in nontraffic accident
V67.3 Unspecified occupant of heavy transport vehicle injured in nontraffic accident
V67.4 Person injured while boarding or alighting
V67.5 Driver injured in traffic accident
V67.6 Passenger injured in traffic accident
V67.7 Person on outside of vehicle injured in traffic accident
V67.9 Unspecified occupant of heavy transport vehicle injured in traffic accident
V68 Occupant of heavy transport vehicle injured in noncollision transport accident
V68.0 Driver injured in nontraffic accident
V68.1 Passenger injured in nontraffic accident
V68.2 Person on outside of vehicle injured in nontraffic accident
V68.3 Unspecified occupant of heavy transport vehicle injured in nontraffic accident
V68.4 Person injured while boarding or alighting
V68.5 Driver injured in traffic accident
V68.6 Passenger injured in traffic accident
V68.7 Person on outside of vehicle injured in traffic accident
V68.9 Unspecified occupant of heavy transport vehicle injured in traffic accident
V69 Occupant of heavy transport vehicle injured in other and unspecified transport accidents
V69.0 Driver injured in collision with other and unspecified motor vehicles in nontraffic accident
V69.1 Passenger injured in collision with other and unspecified motor vehicles in nontraffic accident
V69.2 Unspecified occupant of heavy transport vehicle injured in collision with other and unspecified motor veh
V69.3 Occupant [any] of heavy transport vehicle injured in unspecified nontraffic accident
V69.4 Driver injured in collision with other and unspecified motor vehicles in traffic accident
V69.5 Passenger injured in collision with other and unspecified motor vehicles in traffic accident
V69.6 Unspecified occupant of heavy transport vehicle injured in collision with other and unspecified motor veh
V69.8 Occupant [any] of heavy transport vehicle injured in other specified transport accidents
V69.9 Occupant [any] of heavy transport vehicle injured in unspecified traffic accident
V70 Bus occupant injured in collision with pedestrian or animal
V70.0 Driver injured in nontraffic accident
V70.1 Passenger injured in nontraffic accident
V70.2 Person on outside of vehicle injured in nontraffic accident
V70.3 Unspecified bus occupant injured in nontraffic accident
V70.4 Person injured while boarding or alighting
V70.5 Driver injured in traffic accident
V70.6 Passenger injured in traffic accident
V70.7 Person on outside of vehicle injured in traffic accident
V70.9 Unspecified bus occupant injured in traffic accident

V71 Bus occupant injured in collision with pedal cycle
V71.0 Driver injured in nontraffic accident
V71.1 Passenger injured in nontraffic accident
V71.2 Person on outside of vehicle injured in nontraffic accident
V71.3 Unspecified bus occupant injured in nontraffic accident
V71.4 Person injured while boarding or alighting
V71.5 Driver injured in traffic accident
V71.6 Passenger injured in traffic accident
V71.7 Person on outside of vehicle injured in traffic accident
V71.9 Unspecified bus occupant injured in traffic accident
V72 Bus occupant injured in collision with two- or three-wheeled motor vehicle
V72.0 Driver injured in nontraffic accident
V72.1 Passenger injured in nontraffic accident
V72.2 Person on outside of vehicle injured in nontraffic accident
V72.3 Unspecified bus occupant injured in nontraffic accident
V72.4 Person injured while boarding or alighting
V72.5 Driver injured in traffic accident
V72.6 Passenger injured in traffic accident
V72.7 Person on outside of vehicle injured in traffic accident
V72.9 Unspecified bus occupant injured in traffic accident
V73 Bus occupant injured in collision with car, pick-up truck or van
V73.0 Driver injured in nontraffic accident
V73.1 Passenger injured in nontraffic accident
V73.2 Person on outside of vehicle injured in nontraffic accident
V73.3 Unspecified bus occupant injured in nontraffic accident
V73.4 Person injured while boarding or alighting
V73.5 Driver injured in traffic accident
V73.6 Passenger injured in traffic accident
V73.7 Person on outside of vehicle injured in traffic accident
V73.9 Unspecified bus occupant injured in traffic accident
V74 Bus occupant injured in collision with heavy transport vehicle or bus
V74.0 Driver injured in nontraffic accident
V74.1 Passenger injured in nontraffic accident
V74.2 Person on outside of vehicle injured in nontraffic accident
V74.3 Unspecified bus occupant injured in nontraffic accident
V74.4 Person injured while boarding or alighting
V74.5 Driver injured in traffic accident
V74.6 Passenger injured in traffic accident
V74.7 Person on outside of vehicle injured in traffic accident
V74.9 Unspecified bus occupant injured in traffic accident
V75 Bus occupant injured in collision with railway train or railway vehicle

V75.0 Driver injured in nontraffic accident
V75.1 Passenger injured in nontraffic accident
V75.2 Person on outside of vehicle injured in nontraffic accident
V75.3 Unspecified bus occupant injured in nontraffic accident
V75.4 Person injured while boarding or alighting
V75.5 Driver injured in traffic accident
V75.6 Passenger injured in traffic accident
V75.7 Person on outside of vehicle injured in traffic accident
V75.9 Unspecified bus occupant injured in traffic accident
V76 Bus occupant injured in collision with other nonmotor vehicle
V76.0 Driver injured in nontraffic accident
V76.1 Passenger injured in nontraffic accident
V76.2 Person on outside of vehicle injured in nontraffic accident
V76.3 Unspecified bus occupant injured in nontraffic accident
V76.4 Person injured while boarding or alighting
V76.5 Driver injured in traffic accident
V76.6 Passenger injured in traffic accident
V76.7 Person on outside of vehicle injured in traffic accident
V76.9 Unspecified bus occupant injured in traffic accident
V77 Bus occupant injured in collision with fixed or stationary object
V77.0 Driver injured in nontraffic accident
V77.1 Passenger injured in nontraffic accident
V77.2 Person on outside of vehicle injured in nontraffic accident
V77.3 Unspecified bus occupant injured in nontraffic accident
V77.4 Person injured while boarding or alighting
V77.5 Driver injured in traffic accident
V77.6 Passenger injured in traffic accident
V77.7 Person on outside of vehicle injured in traffic accident
V77.9 Unspecified bus occupant injured in traffic accident
V78 Bus occupant injured in noncollision transport accident
V78.0 Driver injured in nontraffic accident
V78.1 Passenger injured in nontraffic accident
V78.2 Person on outside of vehicle injured in nontraffic accident
V78.3 Unspecified bus occupant injured in nontraffic accident
V78.4 Person injured while boarding or alighting
V78.49 Bus occupant injured in noncollision transport accident; person injured while boarding or alighting; duri
V78.5 Driver injured in traffic accident
V78.6 Passenger injured in traffic accident
V78.7 Person on outside of vehicle injured in traffic accident
V78.9 Unspecified bus occupant injured in traffic accident
V79 Bus occupant injured in other and unspecified transport accidents

V79.0 Driver injured in collision with other and unspecified motor vehicles in nontraffic accident
V79.1 Passenger injured in collision with other and unspecified motor vehicles in nontraffic accident
V79.2 Unspecified bus occupant injured in collision with other and unspecified motor vehicles in nontraffic acci
V79.3 Bus occupant [any] injured in unspecified nontraffic accident
V79.4 Driver injured in collision with other and unspecified motor vehicles in traffic accident
V79.5 Passenger injured in collision with other and unspecified motor vehicles in traffic accident
V79.6 Unspecified bus occupant injured in collision with other and unspecified motor vehicles in traffic acciden
V79.8 Bus occupant [any] injured in other specified transport accidents
V79.89 Bus occupant [any] injured in other specified transport accidents; during unspecified activity
V79.9 Bus occupant [any] injured in unspecified traffic accident
V80 Animal-rider or occupant of animal-drawn vehicle injured in transport accident
V80.0 Rider or occupant injured by fall from or being thrown from animal or animal-drawn vehicle in noncollisi
V80.00 Rider or occupant injured by fall from or being thrown from animal or animal-drawn vehicle in noncollis
V80.01 Rider or occupant injured by fall from or being thrown from animal or animal-drawn vehicle in noncollis
V80.09 Rider or occupant injured by fall from or being thrown from animal or animal-drawn vehicle in noncollis
V80.1 Rider or occupant injured in collision with pedestrian or animal
V80.2 Rider or occupant injured in collision with pedal cycle
V80.3 Rider or occupant injured in collision with two- or three-wheeled motor vehicle
V80.4 Rider or occupant injured in collision with car, pick-up truck, van, heavy transport vehicle or bus
V80.5 Rider or occupant injured in collision with other specified motor vehicle
V80.6 Rider or occupant injured in collision with railway train or railway vehicle
V80.7 Rider or occupant injured in collision with other nonmotor vehicle
V80.8 Rider or occupant injured in collision with fixed or stationary object
V80.9 Rider or occupant injured in other and unspecified transport accidents
V81 Occupant of railway train or railway vehicle injured in transport accident
V81.0 Occupant of railway train or railway vehicle injured in collision with motor vehicle in nontraffic accident
V81.1 Occupant of railway train or railway vehicle injured in collision with motor vehicle in traffic accident
V81.2 Occupant of railway train or railway vehicle injured in collision with or hit by rolling stock
V81.3 Occupant of railway train or railway vehicle injured in collision with other object
V81.4 Person injured while boarding or alighting from railway train or railway vehicle
V81.5 Occupant of railway train or railway vehicle injured by fall in railway train or railway vehicle
V81.6 Occupant of railway train or railway vehicle injured by fall from railway train or railway vehicle
V81.7 Occupant of railway train or railway vehicle injured in derailment without antecedent collision
V81.8 Occupant of railway train or railway vehicle injured in other specified railway accidents
V81.9 Occupant of railway train or railway vehicle injured in unspecified railway accident
V82 Occupant of streetcar injured in transport accident
V82.0 Occupant of streetcar injured in collision with motor vehicle in nontraffic accident
V82.1 Occupant of streetcar injured in collision with motor vehicle in traffic accident
V82.2 Occupant of streetcar injured in collision with or hit by rolling stock
V82.3 Occupant of streetcar injured in collision with other object
V82.4 Person injured while boarding or alighting from streetcar

V82.5 Occupant of streetcar injured by fall in streetcar
V82.6 Occupant of streetcar injured by fall from streetcar
V82.7 Occupant of streetcar injured in derailment without antecedent collision
V82.8 Occupant of streetcar injured in other specified transport accidents
V82.9 Occupant of streetcar injured in unspecified traffic accident
V83 Occupant of special vehicle mainly used on industrial premises injured in transport accident
V83.0 Driver of special industrial vehicle injured in traffic accident
V83.1 Passenger of special industrial vehicle injured in traffic accident
V83.2 Person on outside of special industrial vehicle injured in traffic accident
V83.3 Unspecified occupant of special industrial vehicle injured in traffic accident
V83.4 Person injured while boarding or alighting from special industrial vehicle
V83.5 Driver of special industrial vehicle injured in nontraffic accident
V83.6 Passenger of special industrial vehicle injured in nontraffic accident
V83.7 Person on outside of special industrial vehicle injured in nontraffic accident
V83.72 Person on outside of special industrial vehicle injured in nontraffic accident; while working for income
V83.9 Unspecified occupant of special industrial vehicle injured in nontraffic accident
V84 Occupant of special vehicle mainly used in agriculture injured in transport accident
V84.0 Driver of special agricultural vehicle injured in traffic accident
V84.1 Passenger of special agricultural vehicle injured in traffic accident
V84.2 Person on outside of special agricultural vehicle injured in traffic accident
V84.3 Unspecified occupant of special agricultural vehicle injured in traffic accident
V84.4 Person injured while boarding or alighting from special agricultural vehicle
V84.5 Driver of special agricultural vehicle injured in nontraffic accident
V84.6 Passenger of special agricultural vehicle injured in nontraffic accident
V84.7 Person on outside of special agricultural vehicle injured in nontraffic accident
V84.9 Unspecified occupant of special agricultural vehicle injured in nontraffic accident
V85 Occupant of special construction vehicle injured in transport accident
V85.0 Driver of special construction vehicle injured in traffic accident
V85.1 Passenger of special construction vehicle injured in traffic accident
V85.2 Person on outside of special construction vehicle injured in traffic accident
V85.3 Unspecified occupant of special construction vehicle injured in traffic accident
V85.4 Person injured while boarding or alighting from special construction vehicle
V85.42 Person injured while boarding or alighting from special construction vehicle; while working for income
V85.5 Driver of special construction vehicle injured in nontraffic accident
V85.6 Passenger of special construction vehicle injured in nontraffic accident
V85.7 Person on outside of special construction vehicle injured in nontraffic accident
V85.9 Unspecified occupant of special construction vehicle injured in nontraffic accident
V86 Occupant of special all-terrain or other motor vehicle designed primarily for off-road use, injured in transpo
V86.0 Driver of all-terrain or other off-road motor vehicle injured in traffic accident
V86.1 Passenger of all-terrain or other off-road motor vehicle injured in traffic accident
V86.2 Person on outside of all-terrain or other off-road motor vehicle injured in traffic accident

V86.3 Unspecified occupant of all-terrain or other off-road motor vehicles injured in traffic accident
V86.4 Person injured while boarding or alighting from all-terrain or other off-road motor vehicle
V86.5 Driver of all-terrain or other off-road motor vehicle injured in nontraffic accident
V86.6 Passenger of all-terrain or other off-road motor vehicle injured in nontraffic accident
V86.7 Person on outside of all-terrain or other off-road motor vehicle injured in nontraffic accident
V86.9 Unspecified occupant of all-terrain or other off-road motor vehicle injured in nontraffic accident
V87 Traffic accident of specified type but victim's mode of transport unknown
V87.0 Person injured in collision between car and two- or three-wheeled motor vehicle (traffic)
V87.1 Person injured in collision between other motor vehicle and two- or three-wheeled motor vehicle (traffic)
V87.2 Person injured in collision between car and pick-up truck or van (traffic)
V87.3 Person injured in collision between car and bus (traffic)
V87.4 Person injured in collision between car and heavy transport vehicle (traffic)
V87.5 Person injured in collision between heavy transport vehicle and bus (traffic)
V87.6 Person injured in collision between railway train or railway vehicle and car (traffic)
V87.7 Person injured in collision between other specified motor vehicles (traffic)
V87.8 Person injured in other specified noncollision transport accidents involving motor vehicle (traffic)
V87.9 Person injured in other specified (collision) (noncollision) transport accidents involving nonmotor vehicle
V88 Nontraffic accident of specified type but victim's mode of transport unknown
V88.0 Person injured in collision between car and two- or three-wheeled motor vehicle, nontraffic
V88.1 Person injured in collision between other motor vehicle and two- or three-wheeled motor vehicle, nontraff
V88.2 Person injured in collision between car and pick-up truck or van, nontraffic
V88.3 Person injured in collision between car and bus, nontraffic
V88.4 Person injured in collision between car and heavy transport vehicle, nontraffic
V88.5 Person injured in collision between heavy transport vehicle and bus, nontraffic
V88.6 Person injured in collision between railway train or railway vehicle and car, nontraffic
V88.7 Person injured in collision between other specified motor vehicles, nontraffic
V88.8 Person injured in other specified noncollision transport accidents involving motor vehicle, nontraffic
V88.9 Person injured in other specified (collision) (noncollision) transport accidents involving nonmotor vehicle
V89 Motor- or nonmotor-vehicle accident, type of vehicle unspecified
V89.0 Person injured in unspecified motor-vehicle accident, nontraffic
V89.1 Person injured in unspecified nonmotor-vehicle accident, nontraffic
V89.2 Person injured in unspecified motor-vehicle accident, traffic
V89.3 Person injured in unspecified nonmotor-vehicle accident, traffic
V89.9 Person injured in unspecified vehicle accident
V90 Accident to watercraft causing drowning and submersion
V90.0 Merchant ship
V90.1 Passenger ship
V90.2 Fishing boat
V90.3 Other powered watercraft
V90.4 Sailboat
V90.5 Canoe or kayak

V90.6 Inflatable craft (nonpowered)
V90.7 Water-skis
V90.8 Other unpowered watercraft
V90.9 Unspecified watercraft
V91 Accident to watercraft causing other injury
V91.0 Merchant ship
V91.1 Passenger ship
V91.2 Fishing boat
V91.3 Other powered watercraft
V91.4 Sailboat
V91.5 Canoe or kayak
V91.6 Inflatable craft (nonpowered)
V91.7 Water-skis
V91.8 Other unpowered watercraft
V91.9 Unspecified watercraft
V92 Water-transport-related drowning and submersion without accident to watercraft
V92.0 Merchant ship
V92.1 Passenger ship
V92.2 Fishing boat
V92.3 Other powered watercraft
V92.4 Sailboat
V92.5 Canoe or kayak
V92.6 Inflatable craft (nonpowered)
V92.7 Water-skis
V92.8 Other unpowered watercraft
V92.9 Unspecified watercraft
V93 Accident on board watercraft without accident to watercraft, not causing drowning and submersion
V93.0 Merchant ship
V93.1 Passenger ship
V93.2 Fishing boat
V93.3 Other powered watercraft
V93.31 Accident on board watercraft without accident to watercraft, not causing drowning and submersion; othe
V93.4 Sailboat
V93.5 Canoe or kayak
V93.6 Inflatable craft (nonpowered)
V93.7 Water-skis
V93.8 Other unpowered watercraft
V93.9 Unspecified watercraft
V94 Other and unspecified water transport accidents
V94.0 Merchant ship
V94.1 Passenger ship

V94.2 Fishing boat
V94.3 Other powered watercraft
V94.4 Sailboat
V94.5 Canoe or kayak
V94.6 Inflatable craft (nonpowered)
V94.7 Water-skis
V94.8 Other unpowered watercraft
V94.9 Unspecified watercraft
V95 Accident to powered aircraft causing injury to occupant
V95.0 Helicopter accident injuring occupant
V95.1 Ultralight, microlight or powered-glider accident injuring occupant
V95.2 Accident to other private fixed-wing aircraft, injuring occupant
V95.3 Accident to commercial fixed-wing aircraft, injuring occupant
V95.4 Spacecraft accident injuring occupant
V95.8 Other aircraft accidents injuring occupant
V95.9 Unspecified aircraft accident injuring occupant
V96 Accident to nonpowered aircraft causing injury to occupant
V96.0 Balloon accident injuring occupant
V96.1 Hang-glider accident injuring occupant
V96.2 Glider (nonpowered) accident injuring occupant
V96.8 Other nonpowered-aircraft accidents injuring occupant
V96.9 Unspecified nonpowered-aircraft accident injuring occupant
V97 Other specified air transport accidents
V97.0 Occupant of aircraft injured in other specified air transport accidents
V97.1 Person injured while boarding or alighting from aircraft
V97.2 Parachutist injured in air transport accident
V97.3 Person on ground injured in air transport accident
V97.8 Other air transport accidents, not elsewhere classified
V98 Other specified transport accidents
V99 Unspecified transport accident
W00 Fall on same level involving ice and snow
W00.0 Home
W00.08 Fall on same level involving ice and snow; Home; While engaged in other specified activities
W00.1 Residential institution
W00.2 School, other institution and public administrative area
W00.3 Sports and athletics area
W00.4 Street and highway
W00.42 Fall on same level involving ice and snow; Street and highway; While working for income
W00.5 Trade and service area
W00.6 Industrial and construction area
W00.7 Farm

W00.8 Other specified place
W00.89 Fall on same level involving ice and snow, Other specified places , During unspecified activity
W00.9 Unspecified place
W00.99 Fall on same level involving ice and snow; Unspecified place; During unspecified activity
W01 Fall on same level from slipping, tripping and stumbling
W01.0 Home
W01.03 Fall on same level from slipping, tripping and stumbling; Home; While engaged in other types of work
W01.04 Fall on same level from slipping, tripping and stumbling; Home; While resting, sleeping, eating or engag
W01.08 Fall on same level from slipping, tripping and stumbling; Home; While engaged in other specified activi
W01.09 Fall on same level from slipping, tripping and stumbling; Home; During unspecified activity
W01.1 Residential institution
W01.2 School, other institution and public administrative area
W01.20 Fall on same level from slipping, tripping and stumbling; School, other institution and public administra
W01.22 Fall on same level from slipping, tripping and stumbling; School, other institution and public administra
W01.24 Fall on same level from slipping, tripping and stumbling; School, other institution and public administra
W01.29 Fall on same level from slipping, tripping and stumbling; School, other institution and public administra
W01.3 Sports and athletics area
W01.30 Fall on same level from slipping, tripping and stumbling, Home
W01.31 Fall on same level from slipping, tripping and stumbling; Sports and athletics area; While engaged in lei
W01.4 Street and highway
W01.48 Fall on same level from slipping, tripping and stumbling; Street and highway; While engaged in other sp
W01.49 Fall on same level from slipping, tripping and stumbling; Street and highway; During unspecified activit
W01.5 Trade and service area
W01.53 Fall on same level from slipping, tripping and stumbling; Trade and service area; While engaged in othe
W01.6 Industrial and construction area
W01.7 Farm
W01.8 Other specified place
W01.81 Fall on same level from slipping, tripping and stumbling; Other specified places; While engaged in leisu
W01.88 Fall on same level from slipping, tripping and stumbling; Other specified places; While engaged in other
W01.89 Fall on same level from slipping, tripping and stumbling; Other specified places; During unspecified act
W01.9 Unspecified place
W01.90 Fall on same level from slipping, tripping and stumbling; Unspecified place; While engaged in sports ac
W02 Fall involving ice-skates, skis, roller-skates or skateboards
W02.0 Home
W02.1 Residential institution
W02.2 School, other institution and public administrative area
W02.3 Sports and athletics area
W02.30 Fall involving ice-skates, skis, roller-skates or skateboards; Sports and athletics area; While engaged in s
W02.31 Fall involving ice-skates, skis, roller-skates or skateboards; Sports and athletics area; While engaged in l
W02.4 Street and highway
W02.5 Trade and service area

W02.6 Industrial and construction area
W02.7 Farm
W02.8 Other specified place
W02.81 Fall involving ice-skates, skis, roller-skates or skateboards; Other specified places; While engaged in lei
W02.9 Unspecified place
W03 Other fall on same level due to collision with, or pushing by, another person
W03.0 Home
W03.1 Residential institution
W03.2 School, other institution and public administrative area
W03.3 Sports and athletics area
W03.30 Other fall on same level due to collision with, or pushing by, another person; Sports and athletics area; \
W03.4 Street and highway
W03.5 Trade and service area
W03.6 Industrial and construction area
W03.7 Farm
W03.8 Other specified place
W03.89 Other fall on same level due to collision with, or pushing by, another person, Other specified places , Du
W03.9 Unspecified place
W04 Fall while being carried or supported by other persons
W04.0 Home
W04.1 Residential institution
W04.2 School, other institution and public administrative area
W04.3 Sports and athletics area
W04.4 Street and highway
W04.5 Trade and service area
W04.6 Industrial and construction area
W04.7 Farm
W04.8 Other specified place
W04.9 Unspecified place
W05 Fall involving wheelchair
W05.0 Home
W05.1 Residential institution
W05.2 School, other institution and public administrative area
W05.3 Sports and athletics area
W05.4 Street and highway
W05.5 Trade and service area
W05.6 Industrial and construction area
W05.7 Farm
W05.8 Other specified place
W05.9 Unspecified place
W06 Fall involving bed

W06.0 Home
W06.1 Residential institution
W06.2 School, other institution and public administrative area
W06.28 Fall involving bed; School, other institution and public administrative area; While engaged in other spec
W06.3 Sports and athletics area
W06.4 Street and highway
W06.5 Trade and service area
W06.6 Industrial and construction area
W06.7 Farm
W06.8 Other specified place
W06.9 Unspecified place
W07 Fall involving chair
W07.0 Home
W07.1 Residential institution
W07.2 School, other institution and public administrative area
W07.29 Fall involving chair; School, other institution and public administrative area; During unspecified activity
W07.3 Sports and athletics area
W07.4 Street and highway
W07.5 Trade and service area
W07.6 Industrial and construction area
W07.7 Farm
W07.8 Other specified place
W07.9 Unspecified place
W08 Fall involving other furniture
W08.0 Home
W08.1 Residential institution
W08.2 School, other institution and public administrative area
W08.3 Sports and athletics area
W08.4 Street and highway
W08.5 Trade and service area
W08.6 Industrial and construction area
W08.7 Farm
W08.8 Other specified place
W08.9 Unspecified place
W09 Fall involving playground equipment
W09.0 Home
W09.1 Residential institution
W09.2 School, other institution and public administrative area
W09.3 Sports and athletics area
W09.4 Street and highway
W09.5 Trade and service area

W09.6 Industrial and construction area
W09.7 Farm
W09.8 Other specified place
W09.9 Unspecified place
W10 Fall on and from stairs and steps
W10.0 Home
W10.03 Fall on and from stairs and steps; Home; While engaged in other types of work
W10.04 Fall on and from stairs and steps; Home; While resting, sleeping, eating or engaging in other vital activi
W10.08 Fall on and from stairs and steps; Home; While engaged in other specified activities
W10.1 Residential institution
W10.2 School, other institution and public administrative area
W10.29 Fall on and from stairs and steps; School, other institution and public administrative area; During unspe
W10.3 Sports and athletics area
W10.4 Street and highway
W10.5 Trade and service area
W10.59 Fall on and from stairs and steps; Trade and service area; During unspecified activity
W10.6 Industrial and construction area
W10.7 Farm
W10.8 Other specified place
W10.9 Unspecified place
W10.90 Fall on and from stairs and steps; Unspecified place; While engaged in sports activity
W10.92 Fall on and from stairs and steps; Unspecified place; While working for income
W11 Fall on and from ladder
W11.0 Home
W11.03 Fall on and from ladder; Home; While engaged in other types of work
W11.09 Fall on and from ladder; Home; During unspecified activity
W11.1 Residential institution
W11.2 School, other institution and public administrative area
W11.3 Sports and athletics area
W11.4 Street and highway
W11.5 Trade and service area
W11.6 Industrial and construction area
W11.7 Farm
W11.8 Other specified place
W11.82 Fall on and from ladder; Other specified places; While working for income
W11.9 Unspecified place
W11.99 Fall on and from ladder; Unspecified place; During unspecified activity
W12 Fall on and from scaffolding
W12.0 Home
W12.1 Residential institution
W12.2 School, other institution and public administrative area

W12.3 Sports and athletics area
W12.4 Street and highway
W12.5 Trade and service area
W12.6 Industrial and construction area
W12.62 Fall on and from scaffolding; Industrial and construction area; While working for income
W12.7 Farm
W12.8 Other specified place
W12.9 Unspecified place
W12.92 Fall on and from scaffolding, Unspecified place, While working for income
W12.99 Fall on and from scaffolding; Unspecified place; During unspecified activity
W13 Fall from, out of or through building or structure
W13.0 Home
W13.08 Fall from, out of or through building or structure; Home; While engaged in other specified activities
W13.1 Residential institution
W13.2 School, other institution and public administrative area
W13.3 Sports and athletics area
W13.4 Street and highway
W13.5 Trade and service area
W13.6 Industrial and construction area
W13.7 Farm
W13.8 Other specified place
W13.89 Fall from, out of or through building or structure; Other specified places; During unspecified activity
W13.9 Unspecified place
W14 Fall from tree
W14.0 Home
W14.1 Residential institution
W14.2 School, other institution and public administrative area
W14.3 Sports and athletics area
W14.4 Street and highway
W14.5 Trade and service area
W14.6 Industrial and construction area
W14.7 Farm
W14.8 Other specified place
W14.9 Unspecified place
W15 Fall from cliff
W15.0 Home
W15.1 Residential institution
W15.2 School, other institution and public administrative area
W15.3 Sports and athletics area
W15.4 Street and highway
W15.5 Trade and service area

W15.6 Industrial and construction area
W15.7 Farm
W15.8 Other specified place
W15.9 Unspecified place
W16 Diving or jumping into water causing injury other than drowning or submersion
W16.0 Home
W16.1 Residential institution
W16.2 School, other institution and public administrative area
W16.3 Sports and athletics area
W16.4 Street and highway
W16.5 Trade and service area
W16.6 Industrial and construction area
W16.7 Farm
W16.8 Other specified place
W16.9 Unspecified place
W17 Other fall from one level to another
W17.0 Home
W17.09 Other fall from one level to another, Home , During unspecified activity
W17.1 Residential institution
W17.2 School, other institution and public administrative area
W17.3 Sports and athletics area
W17.4 Street and highway
W17.5 Trade and service area
W17.6 Industrial and construction area
W17.7 Farm
W17.8 Other specified place
W17.81 Other fall from one level to another; Other specified places; While engaged in leisure activity
W17.9 Unspecified place
W17.99 Other fall from one level to another; Unspecified place; During unspecified activity
W18 Other fall on same level
W18.0 Home
W18.04 Other fall on same level, Home , While resting, sleeping, eating or engaging in other vital activities
W18.09 Other fall on same level: Home: During unspecified activity
W18.1 Residential institution
W18.2 School, other institution and public administrative area
W18.22 Other fall on same level; School, other institution and public administrative area; While working for inc
W18.3 Sports and athletics area
W18.4 Street and highway
W18.48 Other fall on same level; Street and highway; While engaged in other specified activities
W18.49 Other fall on same level; Street and highway; During unspecified activity
W18.5 Trade and service area

W18.6 Industrial and construction area
W18.7 Farm
W18.8 Other specified place
W18.89 Other fall on same level; Other specified places; During unspecified activity
W18.9 Unspecified place
W18.93 Other fall on same level; Unspecified place; While engaged in other types of work
W18.99 Other fall on same level; Unspecified place; During unspecified activity
W19 Unspecified fall
W19.0 Home
W19.08 Unspecified fall; Home; While engaged in other specified activities
W19.09 Unspecified fall; Home; During unspecified activity
W19.1 Residential institution
W19.2 School, other institution and public administrative area
W19.3 Sports and athletics area
W19.4 Street and highway
W19.5 Trade and service area
W19.59 Unspecified fall; Trade and service area; During unspecified activity
W19.6 Industrial and construction area
W19.7 Farm
W19.8 Other specified place
W19.81 Unspecified fall; Other specified places; While engaged in leisure activity
W19.9 Unspecified place
W20 Struck by thrown, projected or falling object
W20.0 Home
W20.1 Residential institution
W20.2 School, other institution and public administrative area
W20.22 Struck by thrown, projected or falling object; School, other institution and public administrative area ;W
W20.3 Sports and athletics area
W20.4 Street and highway
W20.49 Struck by thrown, projected or falling object; Street and highway;During unspecified activity
W20.5 Trade and service area
W20.52 Struck by thrown, projected or falling object; Trade and service area;While working for income
W20.6 Industrial and construction area
W20.62 Struck by thrown, projected or falling object; Industrial and construction area ;While working for incom
W20.7 Farm
W20.8 Other specified place
W20.9 Unspecified place
W20.92 Struck by thrown, projected or falling object, Unspecified place, While working for income
W20.99 Struck by thrown, projected or falling object; Unspecified place ;During unspecified activity
W21 Striking against or struck by sports equipment
W21.0 Home

W21.1 Residential institution

W21.2 School, other institution and public administrative area

W21.3 Sports and athletics area

W21.30 Striking against or struck by sports equipment; Sports and athletics area;While engaged in sports activity

W21.4 Street and highway

W21.5 Trade and service area

W21.6 Industrial and construction area

W21.7 Farm

W21.8 Other specified place

W21.9 Unspecified place

W22 Striking against or struck by other objects

W22.0 Home

W22.03 Striking against or struck by other objects; Home; While engaged in other types of work

W22.09 Striking against or struck by other objects; Home; During unspecified activity

W22.1 Residential institution

W22.2 School, other institution and public administrative area

W22.3 Sports and athletics area

W22.4 Street and highway

W22.41 Striking against or struck by other objects; Street and highway;While engaged in leisure activity

W22.5 Trade and service area

W22.6 Industrial and construction area

W22.62 Striking against or struck by other objects; Industrial and construction area ;While working for income

W22.7 Farm

W22.8 Other specified place

W22.88 Striking against or struck by other objects; Other specified places ;While engaged in other specified acti

W22.9 Unspecified place

W23 Caught, crushed, jammed or pinched in or between objects

W23.0 Home

W23.03 Caught, crushed, jammed or pinched in or between objects; Home; While engaged in other types of wor

W23.08 Caught, crushed, jammed or pinched in or between objects; Home; While engaged in other specified act

W23.09 Caught, crushed, jammed or pinched in or between objects; Home; During unspecified activity

W23.1 Residential institution

W23.2 School, other institution and public administrative area

W23.3 Sports and athletics area

W23.4 Street and highway

W23.5 Trade and service area

W23.52 Caught, crushed, jammed or pinched in or between objects; Trade and service area;While working for ir

W23.6 Industrial and construction area

W23.69 Caught, crushed, jammed or pinched in or between objects; Industrial and construction area ;During uns

W23.7 Farm

W23.8 Other specified place

W23.82 Caught, crushed, jammed or pinched in or between objects; Other specified places ;While working for i
W23.9 Unspecified place
W23.92 Caught, crushed, jammed or pinched in or between objects; Unspecified place ;While working for incon
W23.99 Caught, crushed, jammed or pinched in or between objects; Unspecified place ;During unspecified activ
W24 Contact with lifting and transmission devices, not elsewhere classified
W24.0 Home
W24.1 Residential institution
W24.2 School, other institution and public administrative area
W24.3 Sports and athletics area
W24.4 Street and highway
W24.5 Trade and service area
W24.6 Industrial and construction area
W24.7 Farm
W24.8 Other specified place
W24.82 Contact with lifting and transmission devices, not elsewhere classified; Other specified places ;While w
W24.9 Unspecified place
W25 Contact with sharp glass
W25.0 Home
W25.03 Contact with sharp glass; Home; While engaged in other types of work
W25.09 Contact with sharp glass; Home; During unspecified activity
W25.1 Residential institution
W25.18 Contact with sharp glass; Residential institution;While engaged in other specified activities
W25.2 School, other institution and public administrative area
W25.3 Sports and athletics area
W25.4 Street and highway
W25.5 Trade and service area
W25.52 Contact with sharp glass; Trade and service area;While working for income
W25.6 Industrial and construction area
W25.62 Contact with sharp glass; Industrial and construction area ;While working for income
W25.7 Farm
W25.8 Other specified place
W25.9 Unspecified place
W26 Contact with knife, sword or dagger
W26.0 Home
W26.03 Contact with knife, sword or dagger; Home; While engaged in other types of work
W26.08 Contact with knife, sword or dagger, Home , While engaged in other specified activities
W26.09 Contact with knife, sword or dagger; Home; During unspecified activity
W26.1 Residential institution
W26.2 School, other institution and public administrative area
W26.3 Sports and athletics area
W26.4 Street and highway

W26.5 Trade and service area
W26.52 Contact with knife, sword or dagger; Trade and service area;While working for income
W26.6 Industrial and construction area
W26.7 Farm
W26.8 Other specified place
W26.9 Unspecified place
W26.99 Contact with knife, sword or dagger; Unspecified place ;During unspecified activity
W27 Contact with nonpowered hand tool
W27.0 Home
W27.03 Contact with nonpowered hand tool; Home; While engaged in other types of work
W27.09 Contact with nonpowered hand tool; Home; During unspecified activity
W27.1 Residential institution
W27.2 School, other institution and public administrative area
W27.3 Sports and athletics area
W27.4 Street and highway
W27.5 Trade and service area
W27.6 Industrial and construction area
W27.7 Farm
W27.8 Other specified place
W27.9 Unspecified place
W27.92 Contact with nonpowered hand tool; Unspecified place ;While working for income
W27.99 Contact with nonpowered hand tool; Unspecified place ;During unspecified activity
W28 Contact with powered lawn mower
W28.0 Home
W28.00 Contact with powered lawnmower; Home; While engaged in sports activity
W28.01 Contact with powered lawnmower; Home; While engaged in leisure activity
W28.1 Residential institution
W28.2 School, other institution and public administrative area
W28.3 Sports and athletics area
W28.4 Street and highway
W28.5 Trade and service area
W28.6 Industrial and construction area
W28.7 Farm
W28.8 Other specified place
W28.9 Unspecified place
W29 Contact with other powered hand tools and household machinery
W29.0 Home
W29.09 Contact with other powered hand tools and household machinery; Home; During unspecified activity
W29.1 Residential institution
W29.2 School, other institution and public administrative area
W29.3 Sports and athletics area

W29.4 Street and highway
W29.5 Trade and service area
W29.6 Industrial and construction area
W29.62 Contact with other powered hand tools and household machinery; Industrial and construction area ;Whil
W29.7 Farm
W29.8 Other specified place
W29.82 Contact with other powered hand tools and household machinery; Other specified places ;While workin
W29.9 Unspecified place
W30 Contact with agricultural machinery
W30.0 Home
W30.1 Residential institution
W30.2 School, other institution and public administrative area
W30.3 Sports and athletics area
W30.4 Street and highway
W30.5 Trade and service area
W30.6 Industrial and construction area
W30.7 Farm
W30.8 Other specified place
W30.9 Unspecified place
W31 Contact with other and unspecified machinery
W31.0 Home
W31.03 Contact with other and unspecified machinery; Home; While engaged in other types of work
W31.1 Residential institution
W31.2 School, other institution and public administrative area
W31.3 Sports and athletics area
W31.4 Street and highway
W31.5 Trade and service area
W31.52 Contact with other and unspecified machinery; Trade and service area;While working for income
W31.6 Industrial and construction area
W31.62 Contact with other and unspecified machinery; Industrial and construction area ;While working for inco
W31.7 Farm
W31.8 Other specified place
W31.82 Contact with other and unspecified machinery; Other specified places ;While working for income
W31.89 Contact with other and unspecified machinery; Other specified places ;During unspecified activity
W31.9 Unspecified place
W31.92 Contact with other and unspecified machinery; Unspecified place ;While working for income
W32 Handgun discharge
W32.0 Home
W32.1 Residential institution
W32.2 School, other institution and public administrative area
W32.3 Sports and athletics area

W32.4 Street and highway
W32.5 Trade and service area
W32.6 Industrial and construction area
W32.7 Farm
W32.8 Other specified place
W32.9 Unspecified place
W33 Rifle, shotgun and larger firearm discharge
W33.0 Home
W33.1 Residential institution
W33.2 School, other institution and public administrative area
W33.3 Sports and athletics area
W33.4 Street and highway
W33.5 Trade and service area
W33.6 Industrial and construction area
W33.7 Farm
W33.8 Other specified place
W33.9 Unspecified place
W34 Discharge from other and unspecified firearms
W34.0 Home
W34.09 Discharge from other and unspecified firearms; Home; During unspecified activity
W34.1 Residential institution
W34.2 School, other institution and public administrative area
W34.3 Sports and athletics area
W34.4 Street and highway
W34.5 Trade and service area
W34.6 Industrial and construction area
W34.7 Farm
W34.8 Other specified place
W34.9 Unspecified place
W35 Explosion and rupture of boiler
W35.0 Home
W35.1 Residential institution
W35.2 School, other institution and public administrative area
W35.3 Sports and athletics area
W35.4 Street and highway
W35.5 Trade and service area
W35.6 Industrial and construction area
W35.7 Farm
W35.8 Other specified place
W35.9 Unspecified place
W36 Explosion and rupture of gas cylinder

W36.0 Home
W36.1 Residential institution
W36.2 School, other institution and public administrative area
W36.3 Sports and athletics area
W36.4 Street and highway
W36.5 Trade and service area
W36.6 Industrial and construction area
W36.7 Farm
W36.8 Other specified place
W36.9 Unspecified place
W37 Explosion and rupture of pressurised tyre, pipe or hose
W37.0 Home
W37.1 Residential institution
W37.2 School, other institution and public administrative area
W37.3 Sports and athletics area
W37.4 Street and highway
W37.5 Trade and service area
W37.6 Industrial and construction area
W37.7 Farm
W37.8 Other specified place
W37.9 Unspecified place
W38 Explosion and rupture of other specified pressurised devices
W38.0 Home
W38.1 Residential institution
W38.2 School, other institution and public administrative area
W38.3 Sports and athletics area
W38.4 Street and highway
W38.5 Trade and service area
W38.6 Industrial and construction area
W38.7 Farm
W38.8 Other specified place
W38.9 Unspecified place
W39 Discharge of firework
W39.0 Home
W39.1 Residential institution
W39.2 School, other institution and public administrative area
W39.3 Sports and athletics area
W39.4 Street and highway
W39.5 Trade and service area
W39.6 Industrial and construction area
W39.7 Farm

W39.8 Other specified place
W39.9 Unspecified place
W40 Explosion of other materials
W40.0 Home
W40.1 Residential institution
W40.2 School, other institution and public administrative area
W40.3 Sports and athletics area
W40.4 Street and highway
W40.5 Trade and service area
W40.6 Industrial and construction area
W40.7 Farm
W40.8 Other specified place
W40.9 Unspecified place
W41 Exposure to high-pressure jet
W41.0 Home
W41.1 Residential institution
W41.2 School, other institution and public administrative area
W41.3 Sports and athletics area
W41.4 Street and highway
W41.5 Trade and service area
W41.6 Industrial and construction area
W41.7 Farm
W41.8 Other specified place
W41.9 Unspecified place
W42 Exposure to noise
W42.0 Home
W42.1 Residential institution
W42.2 School, other institution and public administrative area
W42.3 Sports and athletics area
W42.4 Street and highway
W42.5 Trade and service area
W42.6 Industrial and construction area
W42.7 Farm
W42.8 Other specified place
W42.9 Unspecified place
W43 Exposure to vibration
W43.0 Home
W43.1 Residential institution
W43.2 School, other institution and public administrative area
W43.3 Sports and athletics area
W43.4 Street and highway

W43.5 Trade and service area
W43.6 Industrial and construction area
W43.7 Farm
W43.8 Other specified place
W43.9 Unspecified place
W44 Foreign body entering into or through eye or natural orifice
W44.0 Home
W44.1 Residential institution
W44.2 School, other institution and public administrative area
W44.3 Sports and athletics area
W44.4 Street and highway
W44.5 Trade and service area
W44.59 Foreign body entering into or through eye or natural orifice; Trade and service area;During unspecified [illegible]
W44.6 Industrial and construction area
W44.7 Farm
W44.8 Other specified place
W44.9 Unspecified place
W44.99 Foreign body entering into or through eye or natural orifice, Unspecified place , During unspecified acti[illegible]
W45 Foreign body or object entering through skin
W45.0 Home
W45.03 Foreign body or object entering through skin; Home; While engaged in other types of work
W45.09 Foreign body or object entering through skin: Home: During unspecified activity
W45.1 Residential institution
W45.2 School, other institution and public administrative area
W45.3 Sports and athletics area
W45.4 Street and highway
W45.5 Trade and service area
W45.6 Industrial and construction area
W45.7 Farm
W45.8 Other specified place
W45.9 Unspecified place
W45.99 Foreign body or object entering through skin; Unspecified place ;During unspecified activity
W46 Contact with hypodermic needle
W46.0 Contact with hypodermic needle (Home)
W46.1 Contact with hypodermic needle (Residential institution)
W46.2 Contact with hypodermic needle (School, other institution and public administrative area)
W46.3 Contact with hypodermic needle (Sports and athletics area)
W46.4 Contact with hypodermic needle (Street and highway)
W46.5 Contact with hypodermic needle (Trade and service area)
W46.6 Contact with hypodermic needle (Industrial and construction area)
W46.7 Contact with hypodermic needle (Farm)

W46.8 Contact with hypodermic needle (Other specified places)
W46.9 Contact with hypodermic needle (Unspecified place)
W49 Exposure to other and unspecified inanimate mechanical forces
W49.0 Home
W49.1 Residential institution
W49.2 School, other institution and public administrative area
W49.3 Sports and athletics area
W49.4 Street and highway
W49.5 Trade and service area
W49.6 Industrial and construction area
W49.7 Farm
W49.8 Other specified place
W49.9 Unspecified place
W50 Hit, struck, kicked, twisted, bitten or scratched by another person
W50.0 Home
W50.1 Residential institution
W50.2 School, other institution and public administrative area
W50.3 Sports and athletics area
W50.30 Hit, struck, kicked, twisted, bitten or scratched by another person; Sports and athletics area;While engag
W50.4 Street and highway
W50.5 Trade and service area
W50.6 Industrial and construction area
W50.7 Farm
W50.8 Other specified place
W50.9 Unspecified place
W51 Striking against or bumped into by another person
W51.0 Home
W51.08 Striking against or bumped into by another person; Home; While engaged in other specified activities
W51.1 Residential institution
W51.2 School, other institution and public administrative area
W51.3 Sports and athletics area
W51.30 Striking against or bumped into by another person: Sports and athletics area: During unspecified activity
W51.4 Street and highway
W51.5 Trade and service area
W51.6 Industrial and construction area
W51.7 Farm
W51.8 Other specified place
W51.81 Striking against or bumped into by another person; Other specified places ;While engaged in leisure acti
W51.9 Unspecified place
W52 Crushed, pushed and stepped on by crowd or human stampede
W52.0 Home

W52.1 Residential institution
W52.2 School, other institution and public administrative area
W52.3 Sports and athletics area
W52.4 Street and highway
W52.5 Trade and service area
W52.6 Industrial and construction area
W52.7 Farm
W52.8 Other specified place
W52.9 Unspecified place
W53 Bitten by rat
W53.0 Home
W53.1 Residential institution
W53.2 School, other institution and public administrative area
W53.3 Sports and athletics area
W53.4 Street and highway
W53.5 Trade and service area
W53.6 Industrial and construction area
W53.7 Farm
W53.8 Other specified place
W53.9 Unspecified place
W54 Bitten or struck by dog
W54.0 Home
W54.08 Bitten or struck by dog; Home; While engaged in other specified activities
W54.09 Bitten or struck by dog; Home; During unspecified activity
W54.1 Residential institution
W54.2 School, other institution and public administrative area
W54.3 Sports and athletics area
W54.4 Street and highway
W54.49 Bitten or struck by dog; Street and highway;During unspecified activity
W54.5 Trade and service area
W54.6 Industrial and construction area
W54.7 Farm
W54.8 Other specified place
W54.81 Bitten or struck by dog; Other specified places ;While engaged in leisure activity
W54.88 Bitten or struck by dog; Other specified places ;While engaged in other specified activities
W54.89 Bitten or struck by dog; Other specified places ;During unspecified activity
W54.9 Unspecified place
W54.93 Bitten or struck by dog; Unspecified place ;While engaged in other types of work
W54.99 Bitten or struck by dog; Unspecified place ;During unspecified activity
W55 Bitten or struck by other mammals
W55.0 Home

W55.1 Residential institution
W55.2 School, other institution and public administrative area
W55.3 Sports and athletics area
W55.4 Street and highway
W55.5 Trade and service area
W55.58 Bitten or struck by other mammals; Trade and service area;While engaged in other specified activities
W55.6 Industrial and construction area
W55.7 Farm
W55.8 Other specified place
W55.9 Unspecified place
W55.99 Bitten or struck by other mammals: Unspecified place: During unspecified activity
W56 Contact with marine animal
W56.0 Home
W56.1 Residential institution
W56.2 School, other institution and public administrative area
W56.3 Sports and athletics area
W56.4 Street and highway
W56.5 Trade and service area
W56.6 Industrial and construction area
W56.7 Farm
W56.8 Other specified place
W56.9 Unspecified place
W57 Bitten or stung by non-venomous insect and other non-venomous arthropods
W57.0 Home
W57.1 Residential institution
W57.2 School, other institution and public administrative area
W57.3 Sports and athletics area
W57.4 Street and highway
W57.5 Trade and service area
W57.6 Industrial and construction area
W57.7 Farm
W57.8 Other specified place
W57.9 Unspecified place
W57.99 Bitten or stung by nonvenomous insect and other nonvenomous arthropods: Unspecified place: During [illegible]
W58 Bitten or struck by crocodile or alligator
W58.0 Home
W58.1 Residential institution
W58.2 School, other institution and public administrative area
W58.3 Sports and athletics area
W58.4 Street and highway
W58.5 Trade and service area

W58.6 Industrial and construction area
W58.7 Farm
W58.8 Other specified place
W58.9 Unspecified place
W59 Bitten or crushed by other reptiles
W59.0 Home
W59.1 Residential institution
W59.2 School, other institution and public administrative area
W59.3 Sports and athletics area
W59.4 Street and highway
W59.5 Trade and service area
W59.6 Industrial and construction area
W59.7 Farm
W59.8 Other specified place
W59.9 Unspecified place
W60 Contact with plant thorns and spines and sharp leaves
W60.0 Home
W60.1 Residential institution
W60.2 School, other institution and public administrative area
W60.3 Sports and athletics area
W60.4 Street and highway
W60.5 Trade and service area
W60.6 Industrial and construction area
W60.7 Farm
W60.8 Other specified place
W60.9 Unspecified place
W64 Exposure to other and unspecified animate mechanical forces
W64.0 Home
W64.1 Residential institution
W64.2 School, other institution and public administrative area
W64.3 Sports and athletics area
W64.4 Street and highway
W64.5 Trade and service area
W64.6 Industrial and construction area
W64.7 Farm
W64.8 Other specified place
W64.9 Unspecified place

tive area; While resting, sleeping, eating or engaging in other vital activities
