## Supplementary Tables 2-5 for "Traumatic Brain Injury Associated with Altered Corpus Callosum Microstructure in Females: Exploring the Roles of Menopause Timing and Hormone Therapy in UK Biobank"

**Part A: Menopause Timing Analysis**

| Region | Measure | F-value | df | p-value | FDR p | Healthy Control mean±SD (n=10128) | TBI Pre-menopause mean±SD (n=30) | TBI Post-menopause mean±SD (n=333) |
|---|---|---|---|---|---|---|---|---|
| Body | FA | 6.411 | 2, 8646 | 0.0016 | **0.0019** | 0.714±0.035 | 0.697±0.051 | 0.710±0.017 |
| Body | MD | 9.9996 | 2, 8646 | <0.0001 | **<0.0001** | 0.00079±0.00005 | 0.00082±0.00007 | 0.00080±0.0 0002 |
| Genu | FA | 8.226 | 2, 8646 | <0.001 | **<0.001** | 0.721±0.040 | 0.706±0.058 | 0.716±0.020 |
| Genu | MD | 8.956 | 2, 8646 | <0.0001 | **<0.0001** | 0.00077±0.00005 | 0.00079±0.00008 | 0.00078±0.0 0003 |
| Genu | ISOVF | 4.031 | 2, 8646 | 0.0178 | **0.0178** | 0.070±0.027 | 0.078±0.040 | 0.073±0.014 |
| Splenium | FA | 7.161 | 2, 8646 | <0.001 | **0.0013** | 0.788±0.026 | 0.775±0.039 | 0.786±0.013 |

**Part B: HRT Effects Analysis**

| Region | Measure | F-value | df | p-value | FDR p | Healthy no HRT mean±SD (n=6436) | Healthy with HRT mean±SD (n=3572) | TBI no HRT mean±SD (n=187) | TBI with HRT mean±SD (n=146) |
|---|---|---|---|---|---|---|---|---|---|
| Body | FA | 8.226 | 3, 8510 | <0.001 | **<0.001** | 0.715±0.042 | 0.712±0.054 | 0.709±0.024 | 0.709±0.025 |
| Body | MD | 14.436 | 3, 8510 | <0.0001 | **<0.0001** | 0.00079±0.00005 | 0.00079±0.00007 | 0.00080±0.00003 | 0.00080±0.00003 |
| Genu | FA | 6.411 | 3, 8510 | 0.0003 | **<0.001** | 0.722±0.048 | 0.718±0.061 | 0.715±0.027 | 0.716±0.029 |
| Genu | MD | 28.041 | 3, 8510 | <0.0001 | **<0.0001** | 0.00077±0.00006 | 0.00077±0.00008 | 0.00078±0.00004 | 0.00078±0.00004 |
| Genu | ISOVF | 28.041 | 3, 8510 | <0.0001 | **<0.0001** | 0.069±0.033 | 0.073±0.042 | 0.073±0.018 | 0.073±0.020 |
| Splenium | FA | 7.161 | 3, 8510 | <0.001 | **<0.001** | 0.789±0.032 | 0.788±0.041 | 0.785±0.018 | 0.786±0.019 |

**Supplementary Table 2, Part A and B.** Corpus callosum microstructure by menopause timing and hormone replacement therapy use**.** Weighted ANOVA results examining group differences in corpus callosum microstructure. Part A compares healthy controls to women with TBI sustained before or after menopause (n=10,128 controls, n=30 pre-menopause TBI, n=333 post-menopause TBI). Part B compares postmenopausal women stratified by TBI status and hormone replacement therapy (HRT) use (n=6,436 healthy no HRT, n=3,572 healthy with HRT, n=187 TBI no HRT, n=146 TBI with HRT). All models adjusted for age at imaging, Townsend deprivation index, educational attainment, and cardiometabolic medication use using doubly robust weighted regression. Group means presented as mean±SD. Abbreviations: FA: fractional anisotropy; MD: mean diffusivity; ISOVF: isotropic volume fraction; FDR: false discovery rate; df: degrees of freedom; TBI: traumatic brain injury; HRT: hormone replacement therapy; SD: standard deviation.

**Part A: Timing Analysis**

| Region | Measure | Contrast | Estimate | SE | t-ratio | p-value | 95% CI Lower | 95% CI Upper | $R^2$ | Adj. $R^2$ |
|---|---|---|---|---|---|---|---|---|---|---|
| Body | FA | Healthy Control - TBI Premenopausal | 0.016559 | 0.0051053 | 3.243 | **0.0034** | 0.004591 | 0.028526 | 0.163 | 0.162 |
| Body | FA | Healthy Control - TBI Postmenopausal | 0.004284 | 0.0017386 | 2.464 | **0.0367** | 0.000208 | 0.008359 | 0.163 | 0.162 |
| Body | FA | TBI Premenopausal - TBI Postmenopausal | -0.012275 | 0.0053789 | -2.282 | 0.0584 | -0.024884 | 0.000334 | 0.163 | 0.162 |
| Body | MD | Healthy Control - TBI Premenopausal | -0.000024 | 0.0000066 | -3.683 | **0.0007** | -0.00004 | -0.000009 | 0.222 | 0.222 |
| Body | MD | Healthy Control - TBI Postmenopausal | -0.000006 | 0.0000022 | -2.567 | **0.0277** | -0.000011 | -0.000001 | 0.222 | 0.222 |
| Body | MD | TBI Premenopausal - TBI Postmenopausal | 0.000018 | 0.0000069 | 2.666 | **0.021** | 0.000002 | 0.000035 | 0.222 | 0.222 |
| Genu | FA | Healthy Control - TBI Premenopausal | 0.014545 | 0.0058308 | 2.494 | **0.0338** | 0.000877 | 0.028212 | 0.214 | 0.213 |
| Genu | FA | Healthy Control - TBI Postmenopausal | 0.005159 | 0.0019857 | 2.598 | **0.0254** | 0.000505 | 0.009814 | 0.214 | 0.213 |
| Genu | FA | TBI Premenopausal - TBI Postmenopausal | -0.009385 | 0.0061433 | -1.528 | 0.2779 | -0.023786 | 0.005015 | 0.214 | 0.213 |
| Genu | ISOVF | Healthy Control - TBI Premenopausal | -0.007952 | 0.0040239 | -1.976 | 0.1182 | -0.017384 | 0.001481 | 0.173 | 0.173 |
| Genu | ISOVF | Healthy Control - TBI Postmenopausal | -0.002817 | 0.0013703 | -2.056 | 0.0993 | -0.006029 | 0.000395 | 0.173 | 0.173 |

| | | | | | | | | | | |
|---|---|---|---|---|---|---|---|---|---|---|
| Genu | ISOVF | TBI Premenopausal - TBI Postmenopausal | 0.005134 | 0.0042395 | 1.211 | 0.4466 | -0.004804 | 0.015072 | 0.173 | 0.173 |
| Genu | MD | Healthy Control - TBI Premenopausal | -0.000023 | 0.0000077 | -3.017 | **0.0072** | -0.000041 | -0.000005 | 0.263 | 0.263 |
| Genu | MD | Healthy Control - TBI Postmenopausal | -0.000008 | 0.0000026 | -2.995 | **0.0078** | -0.000014 | -0.000002 | 0.263 | 0.263 |
| Genu | MD | TBI Premenopausal - TBI Postmenopausal | 0.000015 | 0.0000081 | 1.895 | 0.14 | -0.000004 | 0.000035 | 0.263 | 0.263 |
| Splenium | FA | Healthy Control - TBI Premenopausal | 0.012763 | 0.0038665 | 3.301 | **0.0028** | 0.0037 | 0.021827 | 0.047 | 0.047 |
| Splenium | FA | Healthy Control - TBI Postmenopausal | 0.002475 | 0.0013167 | 1.88 | 0.1447 | -0.000612 | 0.005561 | 0.047 | 0.047 |
| Splenium | FA | TBI Premenopausal - TBI Postmenopausal | -0.010288 | 0.0040738 | -2.526 | **0.0311** | -0.019838 | -0.000739 | 0.047 | 0.047 |

**Part B: HRT Effects Analysis**

| Region | Measure | Contrast | Estimate | SE | t-ratio | p-value | 95% CI Lower | 95% CI Upper | $R^2$ | Adj. $R^2$ |
|---|---|---|---|---|---|---|---|---|---|---|
| Body | FA | Healthy no HRT - Healthy with HRT | 0.003289 | 0.000644 | 5.107 | **<0.0001** | 0.001765 | 0.004813 | 0.168 | 0.168 |
| Body | FA | Healthy no HRT - TBI no HRT | 0.005701 | 0.002379 | 2.396 | **0.0778** | 0.000115 | 0.011287 | 0.168 | 0.168 |
| Body | FA | Healthy no HRT - TBI with HRT | 0.005468 | 0.002544 | 2.149 | 0.1378 | -0.000489 | 0.011425 | 0.168 | 0.168 |
| Body | MD | Healthy no HRT - Healthy with HRT | -0.000005 | 0.00000083 | -6.181 | **<0.0001** | -0.000007 | -0.000003 | 0.226 | 0.226 |
| Body | MD | Healthy no HRT - TBI no HRT | -0.000006 | 0.0000031 | -2.072 | 0.1624 | -0.000013 | 0.000001 | 0.226 | 0.226 |

| | | | | | | | | | | |
|---|---|---|---|---|---|---|---|---|---|---|
| Body | MD | Healthy no HRT - TBI with HRT | -0.000009 | 0.0000033 | -2.846 | **0.023** | -0.000017 | -0.000002 | 0.226 | 0.226 |
| Genu | FA | Healthy no HRT - Healthy with HRT | 0.004123 | 0.000736 | 5.602 | **<0.0001** | 0.002387 | 0.005859 | 0.221 | 0.221 |
| Genu | FA | Healthy no HRT - TBI no HRT | 0.007215 | 0.002719 | 2.654 | **0.0398** | 0.000742 | 0.013688 | 0.221 | 0.221 |
| Genu | FA | Healthy no HRT - TBI with HRT | 0.006319 | 0.002907 | 2.174 | 0.1307 | -0.000595 | 0.013233 | 0.221 | 0.221 |
| Genu | MD | Healthy no HRT - Healthy with HRT | -0.000008 | 0.00000097 | -8.641 | **<0.0001** | -0.000011 | -0.000006 | 0.272 | 0.272 |
| Genu | MD | Healthy no HRT - TBI no HRT | -0.00001 | 0.0000036 | -2.849 | **0.0228** | -0.000018 | -0.000002 | 0.272 | 0.272 |
| Genu | MD | Healthy no HRT - TBI with HRT | -0.000012 | 0.0000038 | -3.15 | **0.0089** | -0.000021 | -0.000003 | 0.272 | 0.272 |
| Genu | ISOVF | Healthy no HRT - Healthy with HRT | -0.004375 | 0.000505 | -8.661 | **<0.0001** | -0.005611 | -0.003139 | 0.182 | 0.182 |
| Genu | ISOVF | Healthy no HRT - TBI no HRT | -0.004227 | 0.001866 | -2.265 | 0.1063 | -0.008799 | 0.000345 | 0.182 | 0.182 |
| Genu | ISOVF | Healthy no HRT - TBI with HRT | -0.004773 | 0.001996 | -2.392 | 0.0787 | -0.009646 | 0.0001 | 0.182 | 0.182 |
| Splenium | FA | Healthy no HRT - Healthy with HRT | 0.001 | 0.000487 | 2.062 | 0.1659 | -0.000188 | 0.002189 | 0.05 | 0.05 |
| Splenium | FA | Healthy no HRT - TBI no HRT | 0.00357 | 0.001801 | 1.982 | 0.1949 | -0.000794 | 0.007934 | 0.05 | 0.05 |
| Splenium | FA | Healthy no HRT - TBI with HRT | 0.00214 | 0.001926 | 1.111 | 0.6825 | -0.00255 | 0.00683 | 0.05 | 0.05 |

**Supplementary Table 3, Part A and B.** Post-hoc pairwise comparisons for significant corpus callosum outcomes. Pairwise comparisons using Tukey's method for outcomes showing significant group differences in Table 3. Part A shows all contrasts for menopause timing analysis (healthy controls, pre-menopause TBI, post-menopause TBI). Part B shows key contrasts for HRT effects analysis; only comparisons with healthy no HRT as reference group are displayed for clarity. Estimates represent differences in DTI metric values between groups from weighted linear models. Positive estimates for FA indicate higher values in the first group; negative estimates for MD and ISOVF indicate lower values in the first group. Models adjusted for age at imaging, Townsend deprivation index, educational attainment, and cardiometabolic medication use. Abbreviations: FA: fractional anisotropy; MD: mean diffusivity; ISOVF: isotropic volume fraction; SE: standard error; CI: confidence interval; TBI: traumatic brain injury; HRT: hormone replacement therapy.

| Variable | Region | Measure | Estimate (95% CI) | P-value | Direction |
|---|---|---|---|---|---|
| Parity | Body | FA | 0.00029 (-0.00018, 0.00077) | 0.231 | Positive |
| Parity | Body | MD | -0.00000 (-0.00000, -0.00000) | **< 0.001** | Negative |
| Parity | Genu | FA | 0.00087 (0.00032, 0.00141) | **0.002** | Positive |
| Parity | Genu | ISOVF | -0.00111 (-0.00149, -0.00074) | **< 0.001** | Negative |
| Parity | Genu | MD | -0.00000 (-0.00000, -0.00000) | **< 0.001** | Negative |
| Parity | Splenium | FA | -0.00121 (-0.00157, -0.00085) | **< 0.001** | Negative |
| Reproductive Span | Body | FA | -0.00004 (-0.00018, 0.00010) | 0.609 | Negative |
| Reproductive Span | Body | MD | 0.00000 (-0.00000, 0.00000) | 0.434 | Positive |
| Reproductive Span | Genu | FA | 0.00017 (0.00001, 0.00033) | **0.037** | Positive |
| Reproductive Span | Genu | ISOVF | -0.00007 (-0.00018, 0.00004) | 0.239 | Negative |
| Reproductive Span | Genu | MD | -0.00000 (-0.00000, 0.00000) | 0.249 | Negative |
| Reproductive Span | Splenium | FA | 0.00007 (-0.00004, 0.00017) | 0.218 | Positive |

**Supplementary Table 4.** Independent associations between reproductive factors and corpus callosum white matter microstructure in menopause timing analysis. Results from linear regression models examining the effects of parity (number of live births) and reproductive span (years from menarche to menopause) on corpus callosum measures among significant outcomes from Analysis 1. Models adjusted for menopause timing group, age at imaging, Townsend deprivation index, educational qualification, cardiometabolic medication use, reproductive span, and parity. Beta coefficients represent the change in outcome per additional year of reproductive span or per additional live birth. Positive estimates for FA indicate higher values with increasing reproductive factor; negative estimates for MD and ISOVF indicate lower values with increasing reproductive factor. Abbreviations: FA: fractional anisotropy; MD: mean diffusivity; ISOVF: isotropic volume fraction; CI: confidence interval.

| Region | Measure | Duration Estimate (95% CI) | Duration P-value | Interaction Beta | Interaction P-value | Age Started Beta | Age Started P-value |
|---|---|---|---|---|---|---|---|
| Body | FA | 0.000009 (-0.000181, 0.000199) | 0.9258 | -0.001079 | 0.0584 | -3.40E-05 | 0.7631 |
| Body | MD | -0.000000 (-0.000000, 0.000000) | **0.0313** | 1e-06 | 0.0637 | 0.00000 | 0.7089 |
| Genu | FA | 0.000084 (-0.000139, 0.000307) | 0.463 | -0.001265 | 0.0577 | 0.000183 | 0.1589 |
| Genu | ISOVF | 0.000089 (-0.000076, 0.000254) | 0.2875 | 0.000771 | 0.1168 | -0.00027 | **0.0049** |
| Genu | MD | -0.000000 (-0.000000, 0.000000) | **0.0145** | 2e-06 | **0.0295** | 0.00000 | 0.0748 |
| Splenium | FA | 0.000530 (0.000383, 0.000677) | **< 0.001** | -0.000402 | 0.3634 | -9.90E-05 | 0.2546 |

**Supplementary Table 5.** HRT duration and timing effects on corpus callosum white matter microstructure among HRT users. Results from linear regression models examining effects of HRT duration (years), TBI×Duration interaction, and age at HRT initiation among women currently or previously using HRT (n=3,108 with valid duration data). Models adjusted for TBI status, age at imaging, Townsend deprivation index, educational qualification, cardiometabolic medication use, and age at menopause. Duration Beta represents the change in outcome per additional year of HRT use. Interaction Beta represents the TBI×Duration interaction term, indicating whether duration effects differ between TBI cases and controls. Age Started Beta represents the effect of age at HRT initiation (per year older at initiation). Positive estimates for FA indicate higher values; positive estimates for MD and ISOVF indicate higher values. Abbreviations: FA: fractional anisotropy; MD: mean diffusivity; ISOVF: isotropic volume fraction; CI: confidence interval.
