## Supplementary figures and images for "Traumatic Brain Injury Associated with Altered Corpus Callosum Microstructure in Females: Exploring the Roles of Menopause Timing and Hormone Therapy in UK Biobank"

### Supplementary Figure 1

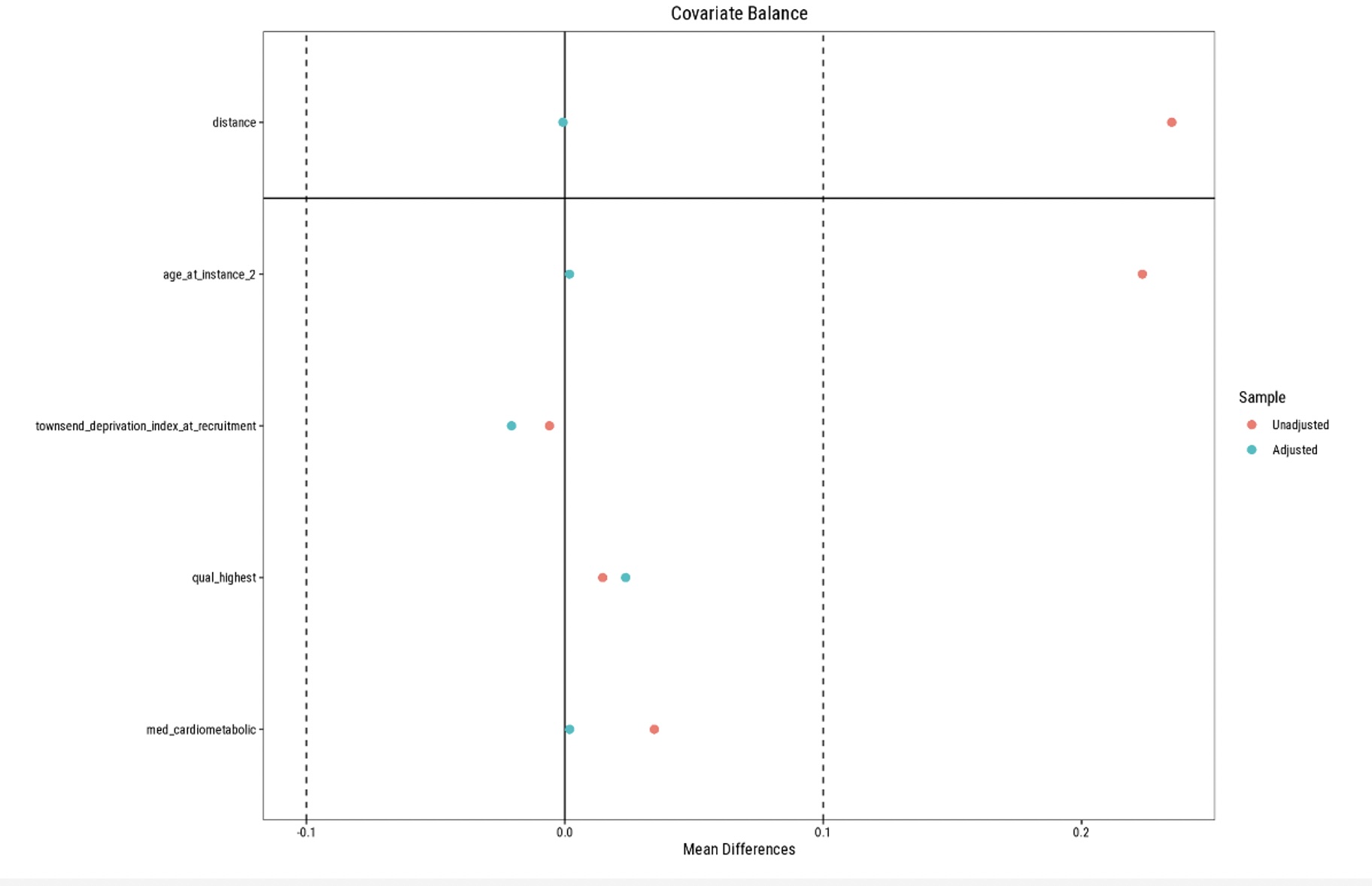
